# HIV-1 Vpr causes separate cell cycle arrests in G2 and M that activate alternative DNA damage pathways

**DOI:** 10.1101/2024.04.02.587683

**Authors:** Ross Hall, Lucy M. Ahern, Melvyn W. Yap, Ming-Han C. Tsai, Virginie C. Boucherit, Tohru Takaki, Simon J. Boulton, Kate N. Bishop

## Abstract

Vpr is a conserved primate lentiviral accessory protein that induces cell cycle arrest in G2. The precise mechanism of this arrest and its benefit to viral replication is unknown. Here, we show that in addition to G2 arrest, Vpr from HIV-1/SIVcpz and HIV-2 lineages separately induce mitotic arrest through the spindle assembly checkpoint, in contrast to other Vpr proteins that only cause G2 arrest. The G2 arrest was mediated solely by ATR (ataxia telangiectasia and Rad3 related) and this activity caused elevated cellular dNTP levels. The mitotic arrest required ATM (ataxia-telangiectasia mutated) as well as ATR activity and resulted from the formation of HIV-1 Vpr-induced ultra-fine anaphase bridges. Moreover, ectopic expression of the DNA structure-specific endonuclease, MUS81, prevented mitotic but not G2 arrest. Importantly, virion-incorporated Vpr was sufficient to induce cellular changes within 12h post-infection, implying that these events benefit the early stages of HIV infection.

**Author Summary:** Vpr is an accessory protein found in primate lentiviruses. Like other retroviral accessory proteins, it is not absolutely required for viral replication but is thought to overcome a cellular factor that negatively regulates infection. The most well-documented effect of Vpr expression is cell cycle arrest in G2. This has been linked to activation of the DNA damage response (DDR) pathway but there are conflicting reports in the literature as to the mechanism behind this. Here, we show that Vpr from some lentiviruses, in fact, cause two separate cell cycle blocks, in G2 and M, that require different DDR pathways. Other Vpr proteins only cause arrest in G2. Furthermore, we show that degradation of one reported target of Vpr, MUS81, is specifically linked to M but not G2 arrest. This indicates that not all Vpr functions are conserved and helps explain contradictory published results. Additionally, we found that virion-incorporated Vpr protein was able to induce cellular changes, including elevated dNTP levels, within 12 hours of infection suggesting that these events enhance early HIV-1 replication events.

## Introduction

Human and simian immunodeficiency viruses (HIV and SIV) encode several accessory proteins that are often dispensable for *in vitro* replication, including viral protein R (Vpr), viral protein U (Vpu), negative regulatory factor (Nef) and viral infectivity factor (Vif) [1]. Some SIV strains and HIV-2, though not HIV-1, encode an additional accessory protein known as Viral protein X (Vpx). Vpr and Vpx have a high sequence similarity, and Vpx is believed to have originated from Vpr by a gene duplication event [2]. Vpr and Vpx are 14-16 kDa proteins and, by virtue of an interaction with the p6 region of Gag, are the only accessory proteins actively packaged into HIV-1 virions [3–6]. Thus, they are present during both early and late stages of infection [7].

Vpx and the Vpr proteins from some SIVs antagonise the cellular protein, Sterile α-motif and histidine-aspartate domain-containing protein 1 (SAMHD1), that restricts viral replication in differentiated cells by reducing dNTP levels to below the threshold required for reverse transcription [8, 9]. However, HIV Vpr does not target SAMHD1. The function of Vpr during HIV infection is currently unknown, although Vpr has been shown to be important for efficient replication in non-dividing macrophages and T lymphocytes [10, 11]. The most well conserved phenotype of Vpr is its ability to induce cell cycle arrest at the G2/M boundary [12–14]. Nonetheless, the cause and purpose for this G2/M arrest is not understood. Vpr-induced cell cycle arrest requires activation of the ataxia telangiectasia and Rad3 related (ATR) kinase, suggesting involvement of the DNA damage response (DDR) pathway [13, 15] although how ATR is activated by Vpr is unclear. Conflicting reports both propose and refute a role for ataxia-telangiectasia mutated (ATM), a protein that is activated in response to double-stranded breaks (DSB) [15, 16]. Occasionally, additional cell cycle defects have been reported, for example expression of Vpr in fission yeast induced numerous defects in the assembly and function of the mitotic spindle and the authors observed similar aberrant mitotic spindles, multiple centrosomes, and multinucleate cells in HeLa cells [17]. A more recent publication identified anaphase-promoting complex, subunit 1 (APC1), a component of the cell cycle master regulator APC/C complex, as a target for Vpr mediated degradation. However, the Vpr protein from NL4-3 did not target APC1, and APC1 degradation did not impact G2 arrest [18].

Vpr binds to DDB1- and CUL4A-associated factor (DCAF1) which is an adaptor for the CRL4/DDB1 E3 ligase complex and this interaction is required for the global proteomic changes in cells that Vpr induces [19], although Vpr also induces widespread transcriptomic changes [20]. Vpr has been reported to target a wide range of cellular substrates for degradation, including, but not limited to, helicase-like transcription factor (HLTF) [21], crossover junction endonuclease MUS81 [22], replication initiation factor MCM10 [23] and uracil-DNA glycosylase (UNG2) [24, 25], all proteins involved in the DDR pathway, and more recently, Tet2 [26], which regulates DNA methylation and CCDC137 [27], which has an unknown function. However, individual knock-down of these proteins does not mimic the Vpr-induced cell cycle arrest. Mutational analysis has shown that the HIV-1 Vpr(Q65R) mutant, which is unable to interact with DCAF1, does not induce cell cycle arrest. In addition, truncation of the C-terminal domain of Vpr, or mutations in this region can also prevent Vpr-induced cell cycle arrest, even though these Vpr proteins can still interact with DCAF1 [28, 29]. Overall, this suggests that Vpr interacts with and degrades a yet unknown cellular substrate, through the C-terminal domain, that results in G2 cell cycle arrest [29–31].

To understand the Vpr-induced cell cycle block further, we studied an extensive panel of Vpr proteins from different HIV/SIV strains and discovered that some Vpr proteins can arrest cells in mitosis in addition to G2. The mitotic block was linked to DNA damage, activation of ATM and degradation of MUS81, although degradation of MUS81 alone was not sufficient. Furthermore, Vpr activation of ATR resulted in elevated dNTP levels within the first 12 hours of infection, when reverse transcription takes place, providing a novel benefit for replication of cell cycle arrest.

## Results

### Some Vpr proteins induce mitotic arrest in addition to arrest in G2

It is well established that Vpr proteins from different strains of HIV/SIV induce cell cycle arrest at the G2/M boundary (**Fig.1A**) [32, 33], although most studies have analysed combined G2 and M populations. To understand this arrest further, we measured the effect of a panel of Vpr proteins on both G2 and M phases. We cloned various lentiviral *vpr* genes into the pLGatewayIRESeYFP retroviral vector and used this to generate MLV transduction vectors encoding FLAG-HA-tagged Vpr proteins upstream of YFP expressed from an internal ribosome entry site (IRES) (**Fig.S1A**). Both Vpr and YFP proteins would be expressed approximately 24 hours post transduction and YFP could then be used as a surrogate for Vpr expression. First, HeLa cells were transduced with vectors encoding either HIV-1_YU2_ Vpr or YU2KO (containing a stop-codon at the start of HIV-1_YU2_ Vpr, preventing expression) or left untransduced (mock). After 48h, cells were fixed, permeabilised and stained with DAPI, to label DNA content, and anti-MPM2(Cy5), which recognizes a variety of proteins that are phosphorylated during mitosis by binding a phospho amino acid-containing epitope (peptides containing LTPLK and FTPLQ sequences) present on more than 50 proteins of M-phase eukaryotic cells, including MAP2, HSP70, cdc25, and DNA topoisomerase IIα, and thus identifies mitotic cells [34]. To avoid the effects of different transduction efficiencies, only YFP-positive cells were analysed by flow cytometry. Cells that had undergone S phase had double the DNA content (4N) and were labelled as G2/M cells. Cells with less DNA content were labelled as G1/S cells. Cells with 4N DNA were then separated into MPM2-positive (mitotic) and negative (G2) populations (**Fig.1B**). Compared to mock cells (**Fig.1B, left panel**), cells transduced with HIV-1_YU2_ Vpr induced a 2.9-fold increase in the percentage of cells in G2 and 4.8-fold increase in the percentage of cells in M phase (**Fig.1B, middle panel**). YU2KO transduction only induced a 1.3-fold, and 1.5-fold increase in the G2 and mitotic populations, respectively (**Fig.1B, right panel**). This suggests that HIV-1_YU2_ Vpr can increase the number of cells in mitosis as well as G2.

**Fig 1.**
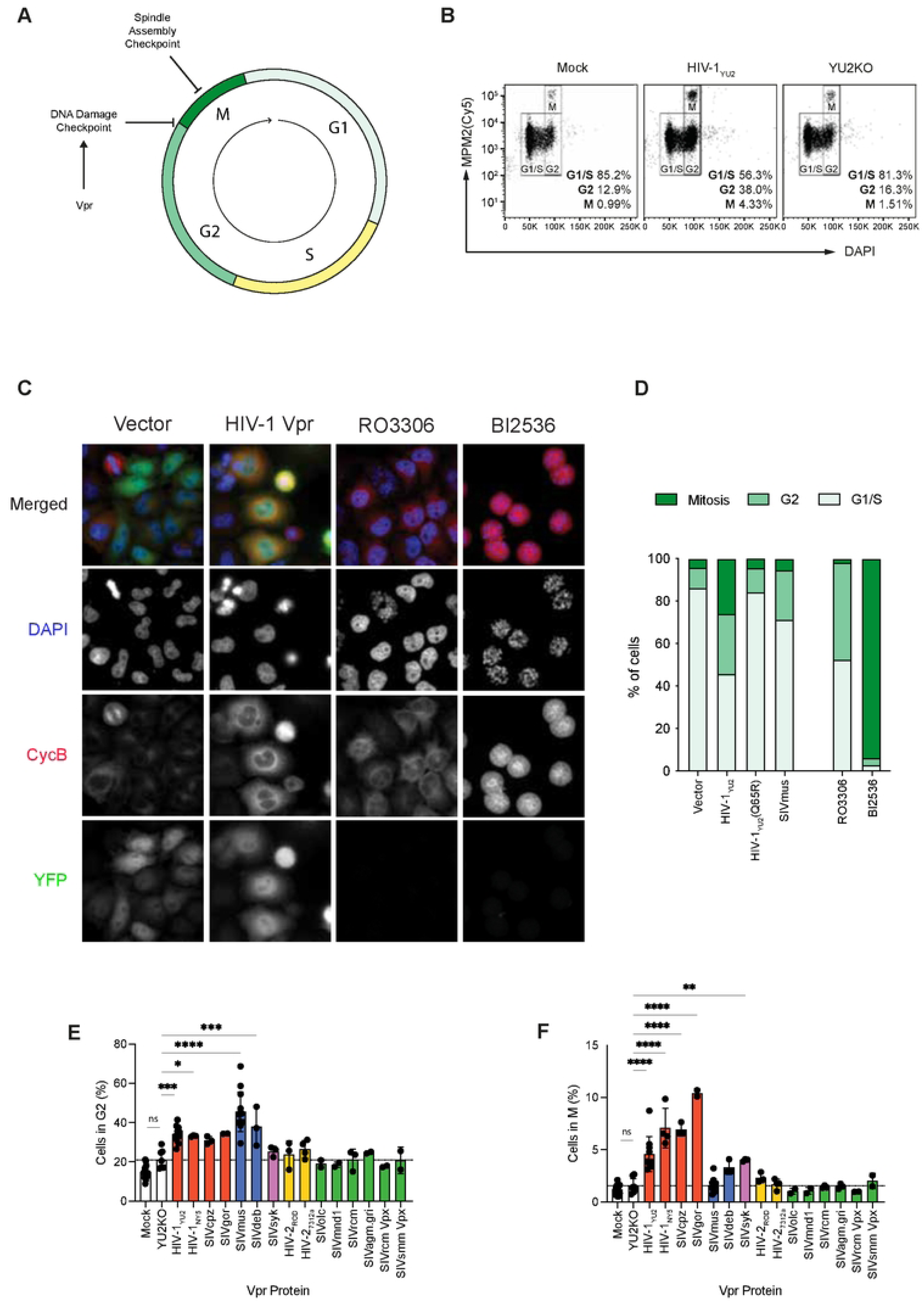
The effect of Vpr on cell cycle arrest. (A) Schematic diagram of the cell cycle and the reported block induced by Vpr. (B) HeLa cells were transduced with MLV transduction vectors encoding FLAG-HA-tagged HIV-1_YU2_ Vpr protein or YU2(KO) upstream of YFP expressed from an IRES (see Fig.S1) or left untransduced (mock). After 48h, cells were harvested, labelled with DAPI and anti-MPM2(Cy5) and the YFP positive cells were analysed by flow cytometry. The gating for different cell cycle phases is shown. (C&D) HeLa cells were transfected with pLGateway-IRES-YFP plasmids encoding HIV-1_YU2_, HIV-1_YU2_(Q65R) or SIV_mus_ Vpr or an empty vector control. Untransduced HeLa cells were treated with PLK1 inhibitor, BI2536, or the selective CDK1 inhibitor, RO-3306 as controls. After 48h, cells were fixed, stained with anti-CycB and DAPI and visualised by confocal microscopy. (C) Representative confocal images. (D) Stack bar plot of the percentage of cells in mitosis, G2 and interphase. (E&F) HeLa cells were transduced with vectors expressing the indicated Vpr protein and cells were analysed by flow cytometry as in (B). The percentage of cells in G2 (E) and M phase (F) was determined. Bar colour groups related lentiviruses. Individual points represent separate biological repeats. Bars show the mean and standard deviation of at least two biological repeats. Statistical analysis was performed using one-way ANOVA to calculate significant differences from the control YU2KO transductions. The significance level was set at * p < 0.05, ** p < 0.01, *** p < 0.001,**** p < 0.0001.

We then used immunofluorescent microscopy to confirm by an alternative method that HIV-1_YU2_ Vpr can arrest cells in both mitosis and G2. HeLa cells were transfected with the pLGatewayIRESeYFP plasmid that expressed HIV-1_YU2_ Vpr, HIV-1_YU2_(Q65R) Vpr with the Q65R mutation that abolishes the interaction with DCAF1 or an empty vector control. Additionally, cells were transfected with a plasmid expressing Vpr from SIV_mus_ which is in a distinct phylogenetic cluster from HIV-1 Vpr and can degrade SAMHD1 [8]. As additional controls, untransduced HeLa cells were treated with the PLK1 inhibitor, BI2536 (to arrest cells in mitosis), or the selective CDK1 inhibitor, RO-3306 (to arrest cells in G2). After 48h, cells were fixed, stained with anti-CycB (to identify cells in G2) and DAPI (to identify condensed mitotic chromatin) and visualised by confocal microscopy **(Fig.1C**). Vpr (YFP) expressing cells were scored as being in mitosis if the chromatin was condensed and in G2 if they were CycB positive but without condensed chromatin. All other cells were labelled as interphase cells. The stack bar plot (**Fig.1D**) shows the percentage of cells in mitosis, G2 and interphase for each Vpr variant. Using this method, HIV-1_YU2_ Vpr expression induced a similar increase in the percentage of cells in both G2 and M compared to the vector control as we detected by flow cytometry. This validates our use of flow cytometry as a more high-throughput and objective method for measuring cell cycle arrest. As expected, the HIV-1_YU2_(Q65R) mutant Vpr failed to increase the percentage of cells in either G2 or M phase (**Fig.1D**). Interestingly, the Vpr from SIV_mus_ only increased the percentage of cells in G2 without affecting the number of cells in M phase (**Fig.1D**).

Therefore, to investigate the conservation of the arrest in mitosis, HeLa cells were transduced with vectors encoding the Vpr protein from a range of lentiviruses: HIV-1_YU2_, HIV-1_NY5_, HIV-2_ROD_, HIV-2_7312a,_ SIV_cpz_, SIV_gor_, SIV_mus_, SIV_deb_, SIV_syk_, SIV_olc_, SIV_mnd1_, SIV_rcm_ or SIV_agm.gri_, or Vpx from SIV_rcm_ and SIV_smm_. Protein expression was confirmed by immunoblotting (**Fig.S1B and C**) and cell cycle arrest was analysed by flow cytometry (**Fig.1E&F**). Compared to the YU2KO control, expression of the Vpr proteins from HIV-1_YU2_, HIV-1_NY5_, SIV_cpz_, SIV_gor_, SIV_mus_ and SIV_deb_ increased the percentage of HeLa cells in G2 (**Fig.1E**). Expression of Vpr from SIV_syk_, HIV-2_ROD_, HIV-2_7312a_ and SIVagm.gri also induced a small increase in the percentage of cells in G2. The remaining Vpr/Vpx proteins did not affect the number of cells in G2. Importantly, cells transduced with Vpr from HIV-1_YU2_, HIV-1_NY5_, SIV_cpz_ and SIV_gor_ also significantly increased the percentage of cells in mitosis (**Fig.1F**), whilst, as in the microscopy experiment, Vpr from SIV_mus_ did not. SIV_deb_ and SIV_syk_ Vpr induced a slight increase in the number of cells in mitosis. No other Vpr/Vpx proteins affected the mitotic population. Altogether, this suggests that blocking the cell cycle in the G2 phase is more prevalent than stalling cells in mitosis, which was mainly limited to Vpr proteins from HIV-1/SIV_cpz_/SIV_gor_ lineage. Therefore, the mitotic block is not just a consequence of blocking cells in G2.

### Mitotic arrest occurs in a range of cell lines

To investigate the breadth of the effect of Vpr on the cell cycle, other human cell lines, U2OS (**Fig.2B**), Jurkat (**Fig.2C**) and SupT1 (**Fig.2D**), and African Green Monkey CV-1 cells (**Fig.2E**) were tested (for complete statistics, see **Fig.S2**). We saw a similar pattern of G2 arrest across all human cell lines, with HIV-1_YU2_, SIV_cpz_, SIV_mus,_ SIV_deb,_ SIV_syk_, HIV-2_ROD_ and SIV_agm.gri_ all increasing the percentage of cells in G2, in some cases to a greater degree than we saw previously in HeLa cells (**Fig.2, left panels**). Furthermore, expression of Vpr from SIV_agm.gri_ induced the strongest G2 block in CV-1 cells (**Fig.2E, left panel**), suggesting that Vpr proteins may be more specific for their cognate host cells, and that although SIVrcm Vpr did not induce a G2 cell cycle block in these lines it may well do so in other monkey cells.

**Fig 2.**
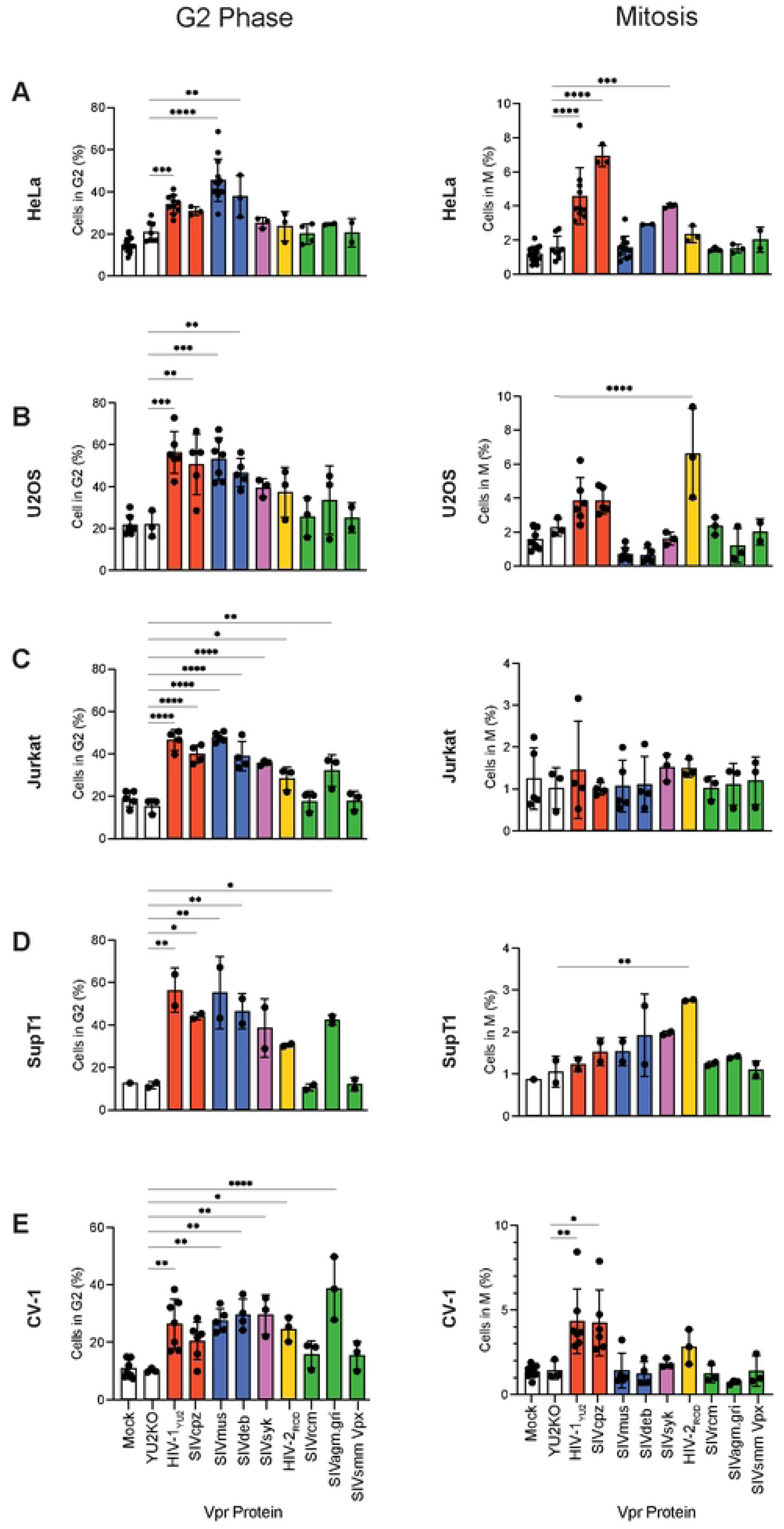
The effect of different Vpr proteins on cell cycle arrest in different cell lines. HeLa (A), U2OS (B), Jurkat (C), SupT1 (D) or CV-1 (E) cells were transduced with MLV transduction vectors encoding the indicated Vpr/Vpx or left untransduced (mock). After 48h, cells were fixed, stained with DAPI and anti-MPM2(Cy5) and the percentage of cells in G2 (left panels) or M phase (right panels) was determined by flow cytometry as in Fig.1. Bar colour groups related lentiviruses. Individual points represent separate biological repeats. Bars show the mean and standard deviation of at least two biological repeats. Statistical analysis was performed using one-way ANOVA to calculate significant differences from the control YU2KO transductions. The significance level was set at * p < 0.05, ** p < 0.01, *** p < 0.001,**** p < 0.0001. See Figure S2 for complete statistical report.

In U2OS and CV-1 cells, HIV-1_YU2_ and SIV_cpz_ Vpr increased the percentage of cells in mitosis as seen in HeLa cells (**Fig.2A, B and E, right panels, red bars**). Interestingly, HIV-2_ROD_ Vpr also increased the percentage of mitotic cells in U2OS and CV-1 cells, despite appearing not to arrest mitosis in HeLa cells (**Fig. 2A, B and E, right panels and Fig.1E, yellow bars**). In these cells, SIV_syk_ had little effect on the M population (**Fig.2B and E, right panels, pink bars**). However, despite causing an increase in the percentage of cells in G2 in the Jurkat and SupT1 T-cell lines, HIV-1_YU2_ and SIV_cpz_ Vpr did not cause a significant increase in the mitotic population in Jurkat and SupT1 cells (**Fig.2C and D**). Surprisingly, HIV-2_ROD_ Vpr did cause an increase in the percentage of mitotic cells in SupT1 cells (**Fig.2D, right panel, yellow bar**). Finally, although expression of SIV_agm.gri_ Vpr increased the number of cells in G2 in CV-1 cells it had no effect on the mitotic population (**Fig.2E**). Thus, differences in blocking various stages of the cell cycle cannot be explained by cell species alone and seems to depend on the exact cell line being tested.

### G2 arrest is different in HIV-1 and SIVmus

We next investigated the timing of the cell cycle arrest induced by Vpr. To accomplish this, we generated doxycycline-inducible stable HeLa cell lines expressing various FLAG-HA-Vpr proteins with YFP expressed from an IRES or only FLAG-HA without Vpr as a control (YFP(only)) (**Fig.S3A).** Stable cell lines were induced with doxycycline for 48h and Vpr expression was detected by immunoblotting (**Fig S3B**). YFP positive cells were sorted (**Fig.S3C**) and further analysed for cell cycle arrest by flow cytometry (**Fig.S3D**). Vpr induction in these stable cells had a similar effect on the percentage of cells in G2 (**Fig.S3D, upper panel**) and M (**Fig.S3D, lower panel**) to that seen in transduced cells (**Fig.1**).

In our experiments, HIV-1_YU2_ Vpr consistently induced the highest levels of G2 arrest of Vpr proteins from the HIV-1/SIV_cpz_/SIV_gor_ lineage (**Fig.1 and Fig.2, red bars**) and induced a strong M phase arrest whereas, although SIV_mus_ Vpr induced the highest levels of G2 arrest of any Vpr protein, it did not arrest cells in M phase (**Fig.1**, **Fig.2 and Fig.S3**). Therefore, in case we were missing an M arrest induced by SIV_mus_ Vpr, we compared how long after induction it took these two Vpr proteins to affect the cell cycle. Cell lines expressing HIV-1_YU2_ or SIV_mus_ Vpr proteins or YFP(only) were induced with doxycycline. Cells were harvested at 16h, 24h, 36h, 48h, 72h and 96h post-induction and the percentages of YFP positive cells in G2 and M phase were determined as before. Cells expressing HIV-1_YU2_ Vpr, had a higher percentage of cells in G2, compared to YFP(only), at all time points (**Fig.3A**), with the peak of G2 arrest seen at the earliest time point measured (16h post-induction). Intriguingly, SIV_mus_ Vpr also increased the percentage of cells in G2, but peaking much later, at 72h (**Fig.3A**). Furthermore, cells expressing HIV-1_YU2_ Vpr also had a higher percentage of cells in mitosis at 16h post-induction, declining after up until 96h (**Fig.3B**), suggesting that HIV-1_YU2_ Vpr causes a rapid inhibition of both G2 and M phases of the cell cycle, likely concomitant with expression, but that SIV_mus_ Vpr causes a slower inhibition of G2 that accumulates with time and does not inhibit mitosis. At the later time points, we noted increased cell death following HIV-1_YU2_ Vpr expression. This may explain the decrease in the number of mitotic cells observed in these cells at 96h post-infection (**Fig.3B**).

**Fig 3.**
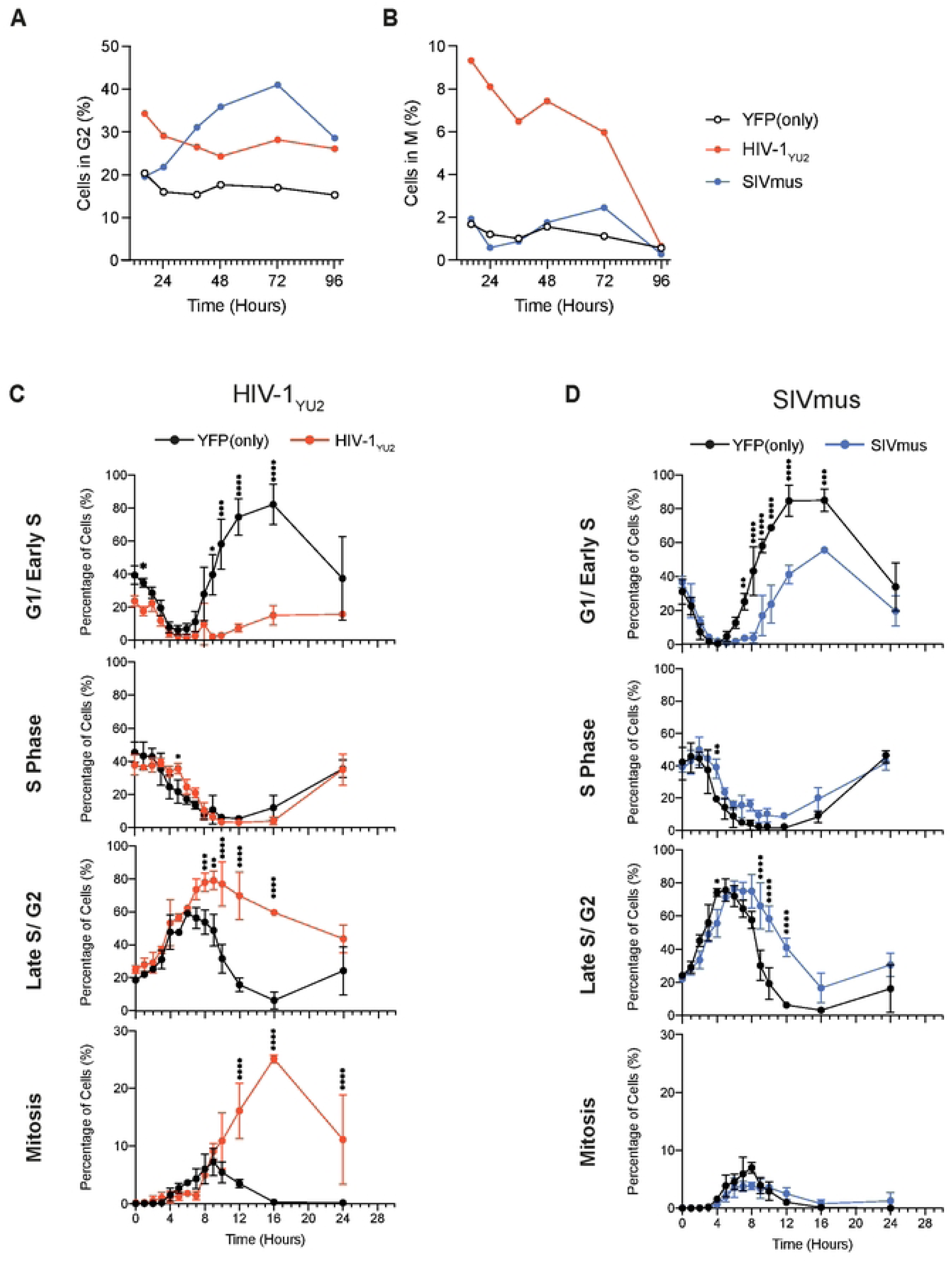
The timing of G2 and M arrest. (A&B) HeLa stable cell lines encoding HIV-1_YU2_, or SIV_mus_ Vpr proteins or YFP(only) were induced with doxycycline and harvested at the indicated timepoint. Cells were labelled with DAPI and anti-MPM2(Cy5) and the percentage of cells in G2 (A) and M phase (B) was determined by flow cytometry. (C&D) HeLa stable cell lines encoding YFP(only) and HIV-1_YU2_ (C) or SIV_mus_ Vpr (D) were induced for 24h. Cells were then treated with Edu (10 μM) for 30 minutes, washed and then harvested at various timepoints (see Fig.S3E). Cells were fixed and immunolabelled with anti-MPM2(Cy5), stained with DAPI and Edu fluorescence was activated. The percentage of YFP positive and Edu positive cells in G1/Early S (1st panel), S phase (2^nd^ panel), Late S/G2 (3^rd^ panel) and mitosis (4^th^ panel) was determined by flow cytometry. Data points represent the mean and standard deviation of two to four separate biological repeats. Statistical analysis was performed using two-way ANOVA to calculate significant differences from the control YFP(only). The significance level was set at * p < 0.05, ** p < 0.01, *** p < 0.001,**** p < 0.0001.

We next tracked cells as they progressed through the cell cycle (S-G2-M-G1) by labelling our stable cell lines with 5-Ethynyl-2’-deoxyuridine (EdU), a thymidine analogue that is incorporated into newly synthesising DNA [35]. This allowed us to follow the mitotic phenotype more closely, which, in asynchronous populations, can be overshadowed by the much larger G2 population. Stable cell lines expressing Vpr from HIV-1_YU2_, SIV_mus_ or the YFP(only) control were induced with doxycycline for 24h before a 30 minute Edu pulse. Doxycycline was maintained in the media for the rest of the experiment. Cells were harvested every hour for ten hours and then at 12h, 16h and 24h (**Fig.S3E**). YFP positive, Edu positive cells were analysed for cell cycle stages by flow cytometry, as outlined in **Fig.S3F**.

**Fig.3C** shows that the percentage of cells in G1/early S decreased until 6h in cells expressing both HIV-1_YU2_ Vpr and YFP(only) (**Fig.3C, top panel**). However, after 6h, the percentage of cells in G1/early S increased in YFP(only) cells, peaking at 16h (**Fig.3C, top panel, black line**), while in HIV-1_YU2_ Vpr expressing cells the percentage of G1/early S cells remained low until the end of the experiment at 24h (**Fig.3C, top panel, red line**), suggesting that Vpr expressing cells were not able to re-enter G1 from mitosis. The percentage of cells in S phase was similar between cells expressing HIV-1_YU2_ Vpr and YFP(only) cells at all time points (**Fig.3C, second panel)**, suggesting that Vpr does not disrupt the G1/S transition or S phase to late S/G2 transition. However, after 6h, the percentage of cells in late S/G2 continued to increase in HIV-1_YU2_ Vpr expressing cells (**Fig.3C, third panel, red line)**, peaking at 9h and then slowly declining until 24h, while in the YFP(only) control cells, the number of cells in Late S/G2 declined from 6h to 16h. This clearly shows the G2 arrest induced by Vpr. Finally, in the YFP(only) cells, the percentage of cells in M phase increased steadily until 9h and then declined (**Fig.3C, bottom panel, black line**) whereas the number of cells in M phase increased more slowly in the HIV-1_YU2_ Vpr expressing cells until 7h, when there was then a more rapid increase until 16h, reaching a level where 25% of the cells were in M phase compared to less that 1% of the control cells (**Fig.3C, bottom panel, red line**). In summary, HIV-1_YU2_ Vpr induced a 3-hour delay in the progression though G2 phase, which reduced the number of mitotic cells that accumulated. However, after this, some cells then proceeded through G2 and further accumulated in M-phase.

SIV_mus_ Vpr expressing cells also showed a delay in cells re-entering G1 compared to the control cells (**Fig.3D, top panel, blue line**) but not as much as HIV-1_YU2_ Vpr expressing cells. SIV_mus_ Vpr did not significantly affect the S phase (**Fig.3D, second panel**). However, as seen for HIV-1 _YU2_ Vpr cells, SIV_mus_ Vpr also caused an increase in cells in Late S/G2 from 6h onwards, compared to control cells (**Fig.3D, third panel**). Notably, the number of cells in G2 declined more rapidly in the SIV_mus_ Vpr lines than in the HIV-1_YU2_ Vpr expressing cells (compare third panels of **Fig.3C and D**), suggesting that the block induced by SIV_mus_ Vpr was not as strong. As seen with HIV-1_YU2_ Vpr, the accumulation of cells in G2 initially reduced the percentage of SIV_mus_ Vpr expressing cells in mitosis compared to the YFP(only) control (**Fig.3D, bottom panel**), but unlike in HIV-1_YU2_ Vpr cells, the percentage of cells in mitosis did not subsequentially increase in SIV_mus_ Vpr expressing cells. This confirms that SIV_mus_ Vpr can delay progression from G2 but does not further inhibit mitosis, and thus does not delay the M/G1 transition as much as HIV-1_YU2_ Vpr. It is interesting to note that while expression of SIV_mus_ Vpr did not appear to induce as strong a G2 arrest as HIV-1_YU2_ Vpr, by percentage of G2 Edu-positive cells (**Fig.3C and D**), it did appear to block successive rounds of replication that resulted in more cells arrested in G2 at 72h compared to HIV-1_YU2_ Vpr (**Fig.3A**). This may be because the strong G2 and M arrests caused by HIV-1_YU2_ Vpr leads to cell death and removal of cells from the total population.

### Mitotic arrest requires DCAF1 and the C-terminus of Vpr

The G2 arrest phenotype is known to be dependent on both the C-terminal region of Vpr and its interaction with DCAF1 [14, 28]. To determine if the same factors were necessary for the block in mitosis, we utilised mutants that knockout the DCAF1 interaction, HIV-1_YU2_(Q65R) and SIV_mus_ Vpr(Q78R), as well as C-terminal truncations and the HIV-1_YU2_(R80A) C-terminal mutant that are thought to be unable to bind to an important Vpr target, failing to bridge it to DCAF, and thereby failing to target it for degradation [14]. HeLa cells were transduced with vectors encoding HIV-1_YU2_, HIV-1_YU2_(Q65R), HIV-1_YU2_(R80A), HIV-1_YU2_(1-79), SIV_mus_, SIV_mus_(Q78R) Vpr proteins, the YU2KO control or left untransduced (mock). After 48h, the percentage of cells in G2 and M phase was determined as before (**Fig.4**). As expected, only the wild type proteins significantly increased the percentage of cells in G2 (**Fig.4A**). Similarly, only expression of wild type HIV-1_YU2_ Vpr increased the percentage of mitotic cells (**Fig.4B, red bar**). Neither SIV_mus_ Vpr nor the SIV_mus_(Q78R) mutant affected mitosis (**Fig.4B**). This suggests that blocking cells in mitosis requires the same interactions with DCAF1 and the C-terminal region of HIV-1 Vpr as are required for the G2 arrest phenotype. To study this further, we generated chimeras between HIV-1_YU2_ and SIV_mus_ Vpr proteins by swapping the C-terminal regions (**Fig.4C**). Removing the C-terminus from either Vpr (HIV-1_YU2_(1-79), SIV_mus_(1-92) and SIV_mus_(1-98)) prevented both from causing a G2 (**Fig.4D**) or M (**Fig.4E**) arrest, as above. However, although replacing the HIV-1_YU2_ C-terminus with the SIV_mus_ C-terminus (HIV-1_YU2_(1-77), SIV_mus_) did not restore G2 arrest (**Fig.4D**), surprisingly, it did restore the M arrest, similar to the wildtype HIV-1_YU2_ Vpr (**Fig.4E**). Replacing the SIV_mus_ C-terminus with the HIV-1_YU2_ C-terminus (SIV_mus_(1-91)HIV-1_YU2_) did not restore either G2 or M arrest. This suggests that the G2 and M arrests can be separated, implying functional differences between these two phenotypes.

**Fig 4.**
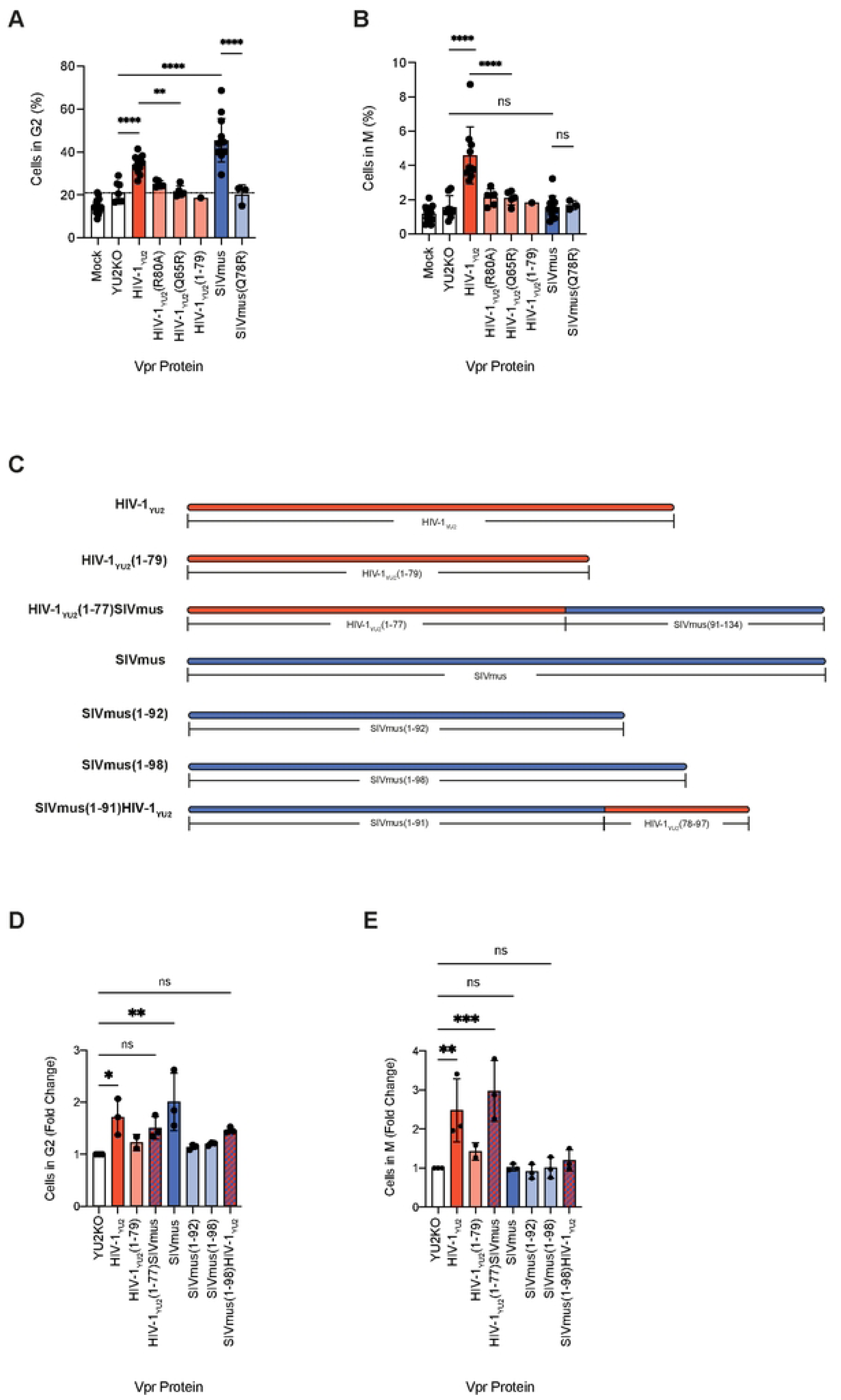
The effect of the C-terminus of Vpr on cell cycle arrest. (A&B) HeLa cells were transduced with vectors encoding HIV-1_YU2_, HIV-1_YU2_(Q65R), HIV-1_YU2_(R80A), HIV-1_YU2_(1-79), SIV_mus_, SIV_mus_(Q78R) Vpr proteins, the YU2KO control or left untransduced (mock). After 48h cells were harvested, labelled with DAPI and anti-MPM2(Cy5) and the percentage of cells in G2 (A) and M phase (B) was determined by flow cytometry. (C) Schematic diagram of HIV-1_YU2_/SIV_mus_ Vpr chimeric proteins indicating the regions corresponding to HIV-1_YU2_ Vpr (red) and SIV_mus_ Vpr (blue). (D&E) HEK293T cells were transfected with plasmids encoding either WT HIV-1_YU2_ or SIV_mus_ Vpr or the chimeric proteins. After 48h, cells were harvested, labelled with DAPI and anti-MPM2(Cy5) and the percentage of cells in G2 and M phase was determined by flow cytometry. The difference in the number of cells in G2 (D) or M phase (E) is reported as fold change compared to the control (YU2KO). Individual points represent separate biological repeats. Bars show the mean and standard deviation of at least three biological repeats. Statistical analysis was performed using one-way ANOVA to calculate significant differences from the control YU2KO transductions. The significance level was set at * p < 0.05, ** p < 0.01, *** p < 0.001,**** p < 0.0001. ns indicates not significant.

### HIV-1 causes DNA damage and induces spindle assembly defects

HIV-1 Vpr expression increased the percentage of cells in mitosis, suggesting activation of the mitotic cell cycle checkpoint. A delay in mitosis is usually activated in response to chromosome misalignment and segregation stress [36]. To determine if HIV-1_YU2_ Vpr induced chromosome misalignment, HeLa cells were transfected with pLGateway-IRES-YFP plasmids encoding HIV-1_YU2_, HIV-1_YU2_(Q65R) or SIV_mus_ Vpr or an empty vector control. After 48h, cells were fixed, stained with anti-tubulin and DAPI and visualised by confocal microscopy. YFP (Vpr) positive cells clearly showed chromosomal abnormalities, indicative of failed mitosis due to misalignment and segregation of condensed chromosomes (**Fig.5A and Fig.S4A&B**). Therefore, we investigated whether we could detect DNA damage during the mitotic phase. Our HeLa stable cell lines were induced to express HIV-1_YU2_, HIV-1_YU2_(R80A), HIV-1_YU2_(Q65R) or SIV_mus_ Vpr or YFP(only), and stained with anti-pH2AX as a DNA damage marker [37], anti-MPM2(Cy5) and DAPI. The percentage of cells with phosphorylated H2AX (γH2AX) in different phases of the cell cycle was then determined by flow cytometry (**Fig.5B**). As a positive control, cells were treated with hydroxyurea (HU) for 8h before staining. This resulted in increased γH2AX staining in cells in G1/S and G2 but not in cells in mitosis. The percentage of γH2AX positive cells was lower than seen in microscopy experiments suggesting that flow cytometry is a less sensitive method of detecting γH2AX foci, or that our HU treatment was sub-optimal. Strikingly however, cells expressing HIV-1_YU2_ Vpr showed an increase in the γH2AX signal in all cell cycle phases, particularly in G2 and M phases, compared to cells expressing the Vpr mutants and YFP(only), indicating that Vpr expression causes DNA damage but in a different way to hydroxyurea. Consistent with the lack of M arrest, SIV_mus_ showed no significant change in γH2AX signal in mitotic cells compared to YFP(only) (**Fig.5B**).

**Fig 5.**
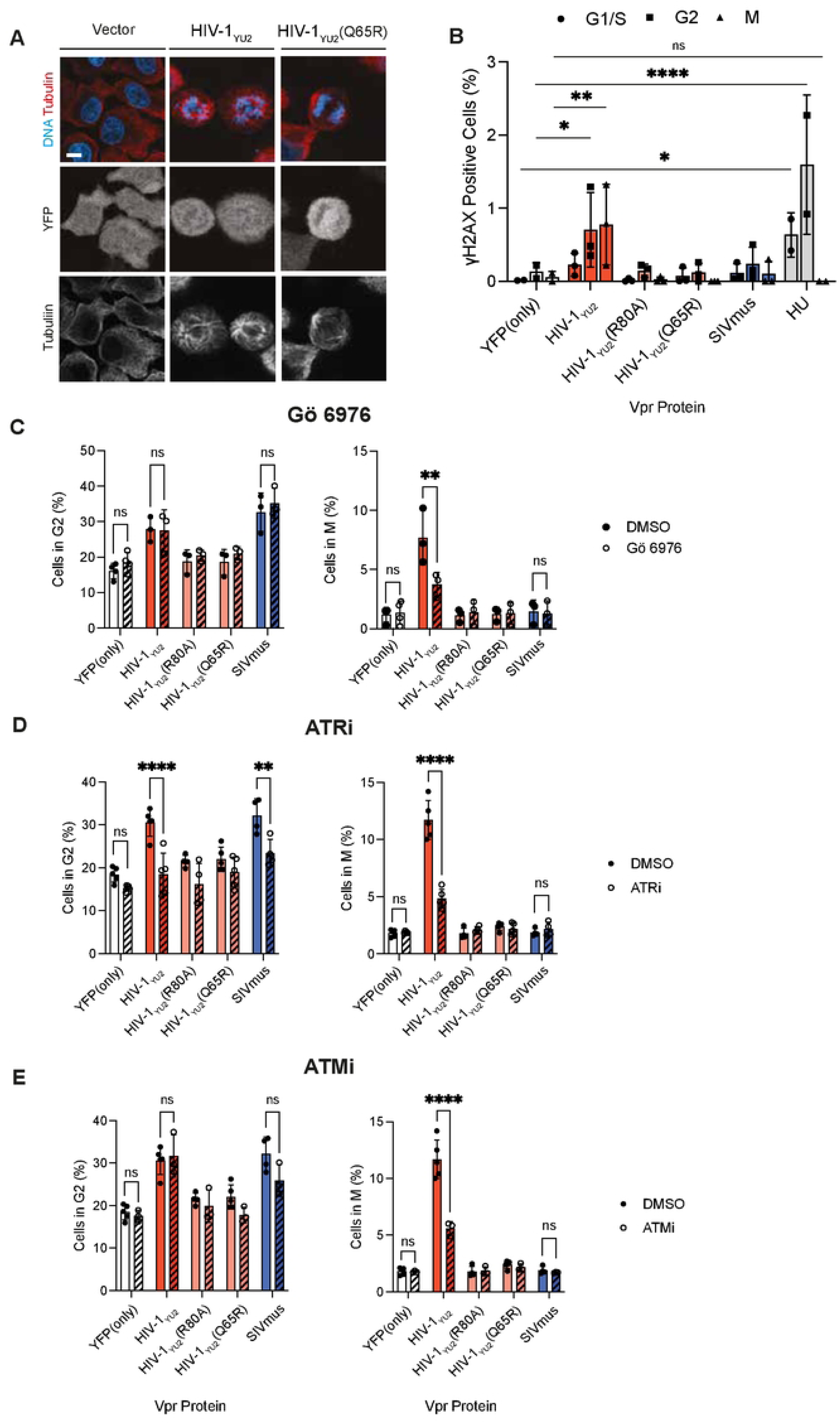
The role of the spindle assembly checkpoint, ATR and ATM kinases in Vpr function. (A) HeLa cells were transfected with pLRGateway2IRESeYFP plasmids encoding HIV-1_YU2_ or HIV-1_YU2_(Q65R) Vpr or an empty vector control. After 48h, cells were fixed, stained with anti-tubulin and DAPI and visualised by confocal microscopy. (B) HeLa stable cell lines encoding HIV-1_YU2_, HIV-1_YU2_(R80A), HIV-1_YU2_(Q65R) or SIV_mus_ Vpr or YFP(only) were induced with doxycycline for 24h. As a control, HeLa cells were treated with 200mM HU for 8h. Cells were then fixed and stained with anti-γH2AX(PE), anti-MPM2(Cy5) and DAPI. The percentage of γH2AX positive cells in each phase of the cell cycle was determined by flow cytometry. Individual points represent separate biological repeats. Statistical analysis was performed using two-way ANOVA to calculate significant differences from the control YFP(only) for each cell cycle stage. (C) HeLa stable cell lines encoding the indicated Vpr were induced with doxycycline for 22h. Cells were then treated with the PKC inhibitor, Gö 6976, as well as doxycycline for a further 2h before staining with anti-MPM2(Cy5) and DAPI. The percentage of cells in G2 and M phase was analysed by flow cytometry. (D&E) HeLa stable cell lines encoding the indicated Vpr were induced with doxycycline for 24h before cells were treated with either an inhibitor of ATR (VE-821) or ATM (KU-55933) for 24h. The percentage of cells in G2 and M phases was measured as in (C). (C-E) Individual points represent separate biological repeats. Bars show the mean and standard deviation of at least three biological repeats. Statistical analysis was performed using two-way ANOVA to calculate significant differences from the DMSO control. The significance level was set at * p < 0.05, ** p < 0.01, *** p < 0.001,**** p < 0.0001. ns indicates not significant.

As HIV-1_YU2_ Vpr induced both DNA damage and spindle abnormalities, we examined whether the spindle assembly checkpoint (SAC), was activated in response to Vpr expression. The SAC activates the mitotic checkpoint complex (MCC), composed of MAD2, Cdc20, BubR1 and BUB3. This complex prevents APC/C-dependent degradation of Cyclin B1, delaying mitotic exit to give time for DNA damage repair and/or chromosome alignment to occur [38, 39]. Firstly, we induced our stable Vpr HeLa cell lines with doxycycline for 22h and then treated the cells with the PKC inhibitor, Gö6976, which inhibits the SAC by targeting Aurora-A and B kinases [40]. After 2h, cells were stained with anti-MPM2(Cy5) and DAPI and analysed by flow cytometry, **Fig.5C**. As expected, treatment of cells with Gö6976 had no significant effect on the G2 populations (**Fig.5C, left panel**). However, it did cause a decrease in the percentage of HIV-1_YU2_ Vpr expressing cells in M phase, compared to the DMSO control (**Fig.5C, right panel**). Gö6976 had no effect on mitotic cell population for YFP(only), HIV-1_YU2_(R80A) and HIV-1_YU2_(Q65R) or SIV_mus_ expressing cells (**Fig.5C, right panel**). This suggests that HIV-1_YU2_ Vpr activates the SAC resulting in mitotic delay. A similar result was observed when co-expressing a dominant negative form of MAD2 (MAD2-3SD) [41] (**Fig.S4C**).

### HIV-1 induces mitotic arrest through ATR and ATM kinases

Previous work has shown that Vpr activates the ATR DNA damage response to induce G2 arrest [13, 42]. Whether the ATM pathway is involved is more controversial [15, 16]. To investigate the role of ATM and ATR in Vpr-induced cell cycle delay, Vpr was induced in our stable HeLa cells before cells were treated with potent and selective inhibitors of ATR (VE-821) or ATM (KU-55933). The percentage of cells in G2 and M phases was measured as above. As previously reported, ATR inhibition prevented the G2 arrest induced by HIV-1_YU2_ and SIV_mus_ Vpr (**Fig.5D, left panel**), while ATM inhibition had no effect on the G2 populations (**Fig.5E, left panel**). Interestingly, the increase in the number of cells in mitosis induced by HIV-1_YU2_ Vpr was abrogated by inhibition of ATR (**Fig.5D, right panel**) and ATM (**Fig.5E, right panel**). ATM inhibition did not change the percentage of cells in G2 or mitosis in any of the other cell lines. This suggests that HIV-1_YU2_ Vpr can activate both ATR and ATM signalling, although G2 arrest only requires ATR activity.

### Expression of HIV-1_YU2_ Vpr induces ultra-fine anaphase bridges

Vpr has been reported to increase FANCD2 foci in cells, suggesting the presence of anaphase bridges [22]. Anaphase bridges are ultra-fine DNA strands that arise from either incomplete replication during S phase or incomplete homologous recombination in response to DNA damage [43]. If bridges occur during mitosis, they can prevent chromosome segregation and delay mitotic exit. Thus, we investigated whether Vpr induced anaphase bridges and if this correlated with the ability of Vpr to inhibit mitosis. HEK293T cells were transfected with plasmids expressing either HIV-1_YU2_, HIV-1_YU2_(R80A), HIV-1_YU2_(Q65R), YU2KO or SIV_mus_ Vpr, stained with anti-replication protein A (RPA, for anaphase bridges), and DAPI and analysed by confocal microscopy (**Fig.6A**). HIV-1_YU2_ Vpr expression significantly increased the number of anaphase bridges during mitosis as seen by RPA staining (**Fig.6A and B**) compared to the YU2KO control, while SIV_mus_ Vpr expression did not (**Fig.6B**). The HIV-1_YU2_(R80A) also induced a small increase in the number of anaphase bridges (**Fig.6B**). We were unable to observe FANCD2 foci flanking the anaphase bridges in the presence of HIV-1_YU2_ (**Fig.6A and C**), suggesting that the bridges arise from incomplete homologous recombination rather than replication stress [43].

**Fig 6.**
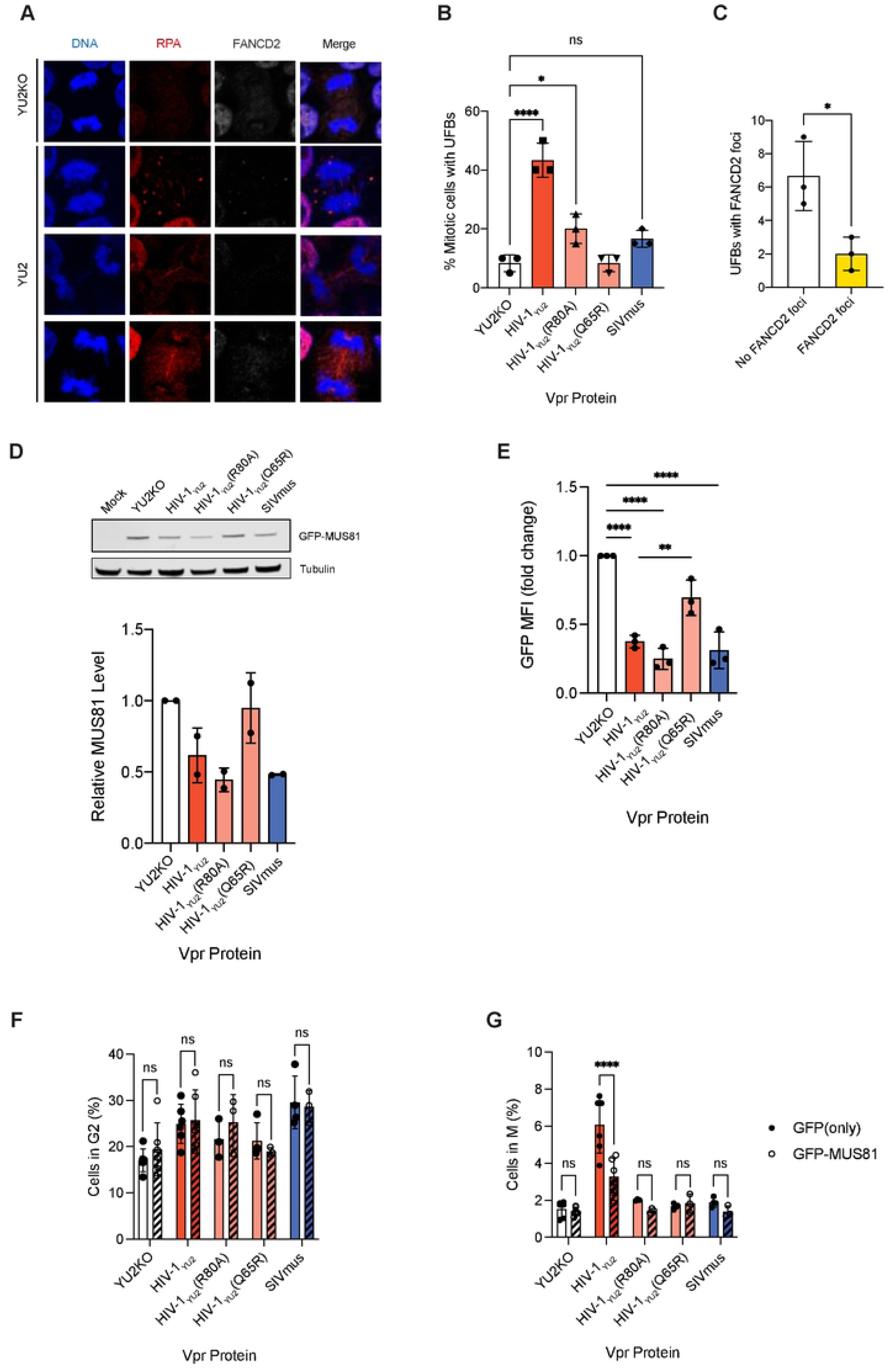
The effect of Vpr on ultra-fine anaphase bridges and MUS81. HEK293T cells were transfected with pLRGateway2IRESeYFP plasmids expressing either HIV-1_YU2_, HIV-1_YU2_(R80A), HIV-1_YU2_(Q65R), SIV_mus_ Vpr or YU2KO. (A-C) After 48h cells were fixed and labelled with anti-RPA, anti-FANCD2 and DAPI and analysed by confocal microscopy (A). The percentage of mitotic cells with RPA labelled ultra-fine bridges (UFB) was quantified (B) and individual UFB were analysed for the presence of FANCD2 foci at their termini (C). Individual points represent separate biological repeats. Bars show the mean and standard deviation of at least three biological repeats. Statistical analysis was performed using an unpaired t-test. (D-G) Cells were simultaneously co-transfected with plasmids encoding GFP-MUS81 or GFP(only). After 24h, cells were harvested and analysed: (D) Cells were lysed and analysed by immunoblotting for anti-GFP and anti-tubulin. Graph shows quantification of MUS81 levels, normalised to tubulin, relative to cells expressing YU2KO. (E) Cells were analysed by flow cytometry for GFP mean fluorescence intensity (MFI). The fold change in intensity relative to cells expressing YU2KO is plotted. Statistical analysis was performed using two-way ANOVA to calculate significant differences from the control YU2KO transfection. (F&G) Cells were fixed, stained with anti-MPM2(Cy5) and DAPI and the percentage of cells in G2 (F) and M phase (G) was analysed by flow cytometry. Individual points represent separate biological repeats. Bars show the mean and standard deviation of at least three biological repeats. Statistical analysis was performed using two-way ANOVA to calculate significant differences from the GFP(only) control. The significance level was set at * p < 0.05, ** p < 0.01, *** p < 0.001,**** p < 0.0001. ns indicates not significant.

### Ectopic expression of MUS81 can rescue mitotic arrest

HIV-1 Vpr can target the SLX4-MUS81 complex for degradation [22, 44]. MUS81 is a structure-specific endonuclease that is involved in resolving anaphase bridges [43, 45]. We therefore examined whether MUS81 degradation is required for the cell cycle arrest phenotypes. First, we ectopically expressed GFP-MUS81 in cells (**Fig.S5A**) and confirmed that Vpr could degrade GFP-MUS81 by western blotting (**Fig.6D**) and flow cytometry (**Fig.6E**). Interestingly, in addition to HIV-1_YU2_ and SIV_mus_ Vpr, the HIV-1_YU2_(R80A) mutant could also degrade GFP-MUS81, which did not correlate with ability to cause either G2 or M arrest. As shown previously [22], HIV-1_YU2_(Q65R) could not degrade MUS81, affirming that interaction with DCAF1 is required for this activity (**Fig.6D and E**). Next, we determined if ectopic expression of GFP-MUS81, could reduce cell cycle arrest. HEK293T cells were transfected with plasmids expressing either HIV-1_YU2_, HIV-1_YU2_(R80A), HIV-1_YU2_(Q65R), YU2KO or SIV_mus_ Vpr, as well as a plasmid encoding GFP-MUS81 or just GFP as a control and analysed by flow cytometry to determine the percentage of cells in G2 and M phases. Co-expression of GFP or GFP-MUS81 had no effect on the ability of HIV-1_YU2_ or SIV_mus_ Vpr to induce G2 arrest (**Fig.6F**). However, expressing GFP-MUS81 significantly reduced the percentage of cells in mitosis compared to the GFP control in cells expressing HIV-1_YU2_ Vpr (**Fig.6G**). Mitosis was unaffected by GFP-MUS81 expression in cells expressing the other Vpr proteins. We also tested whether ectopic expression of other structure-specific endonucleases, XPF and GEN1, could abrogate the mitotic block. While expression of XPF-GFP reduced the percentage of cells in mitosis in HIV-1_YU2_ Vpr expressing cells (**Fig.S5B, right panel**), ectopic expression of GEN1 had no effect on the ability of HIV-1_YU2_ Vpr to cause G2 or M arrest (**Fig.S5C**). We also tested the effect of ectopically expressing another reported target of Vpr, CCDC137 [27] which has no known function (**Fig.S5D**). Expressing CCDC137 did not rescue the mitotic arrest induced by HIV-1_YU2_ Vpr. Instead, ectopic expression of CCDC137 appeared to augment both G2 and M arrest in both HIV-1_YU2_ Vpr expressing cells and control cells (**Fig.S5D**).

### Vpr causes increases in cellular dNTP levels

The advantage of inducing cell cycle arrest for viral replication is unknown. One reasonable suggestion is that inducing DNA damage or arresting the cell cycle may alter the availability of dNTPs, which are known to fluctuate during the cell cycle and in response to DNA damage. Altering dNTP pools by degrading the deoxynucleoside triphosphohydrolase, SAMHD1, is a known function of Vpx and some Vpr proteins [22, 46, 47]. Therefore, HeLa stable Vpr cell lines were induced with doxycycline and dNTP levels were determined using a primer extension assay [9, 46]. Compared to cells expressing YFP(only), cells expressing HIV-1_YU2_ Vpr had increased levels of all four dNTPs (**Fig.7A**), although only the increase in dTTP was significant. Interestingly, expression of both HIV-1_NY5_ and SIV_mus_ Vpr proteins also increased dNTP levels (**Fig.7B, red and blue bars**) whereas HIV-2_ROD_ Vpr did not (**Fig.7B, yellow bar**). This correlated to the G2 arrest rather than the M arrest that was only induced by HIV-1 Vpr proteins in HeLa cells (**Fig.1**). To investigate this, we treated the stable HeLa cells with increasing concentrations of ATR inhibitor and compared the effect on G2/M arrest to the effect on dATP and dTTP levels. As expected from **Fig.5**, treatment with the ATR inhibitor resulted in a dose dependent decrease in G2/M arrest induced by HIV-1_YU2_ Vpr (**Fig.7C**). It also reduced dNTP levels back to the levels in control cells (**Fig.7D**). To study this further, we made stable Jurkat T-cell lines expressing the doxycycline inducible Vpr constructs (**Fig.S3A**) as HIV-1_YU2_ Vpr expression induced a G2 arrest but not an M arrest in Jurkat cells (**Fig.2**). In stable, Vpr-inducible, Jurkat cells, induction of HIV-1_YU2_ Vpr also increased dNTP levels similar to HeLa cells (**Fig.7E**). Furthermore, the dNTP level rise was abrogated by the R80A and Q65R Vpr mutations (**Fig.7E**) showing that it is dependent on the C-terminal region of Vpr and DCAF1 interaction, just like the cell cycle blocks. Altogether, this suggests that increasing dNTP levels is linked to the Vpr-induced G2 arrest.

**Fig 7.**
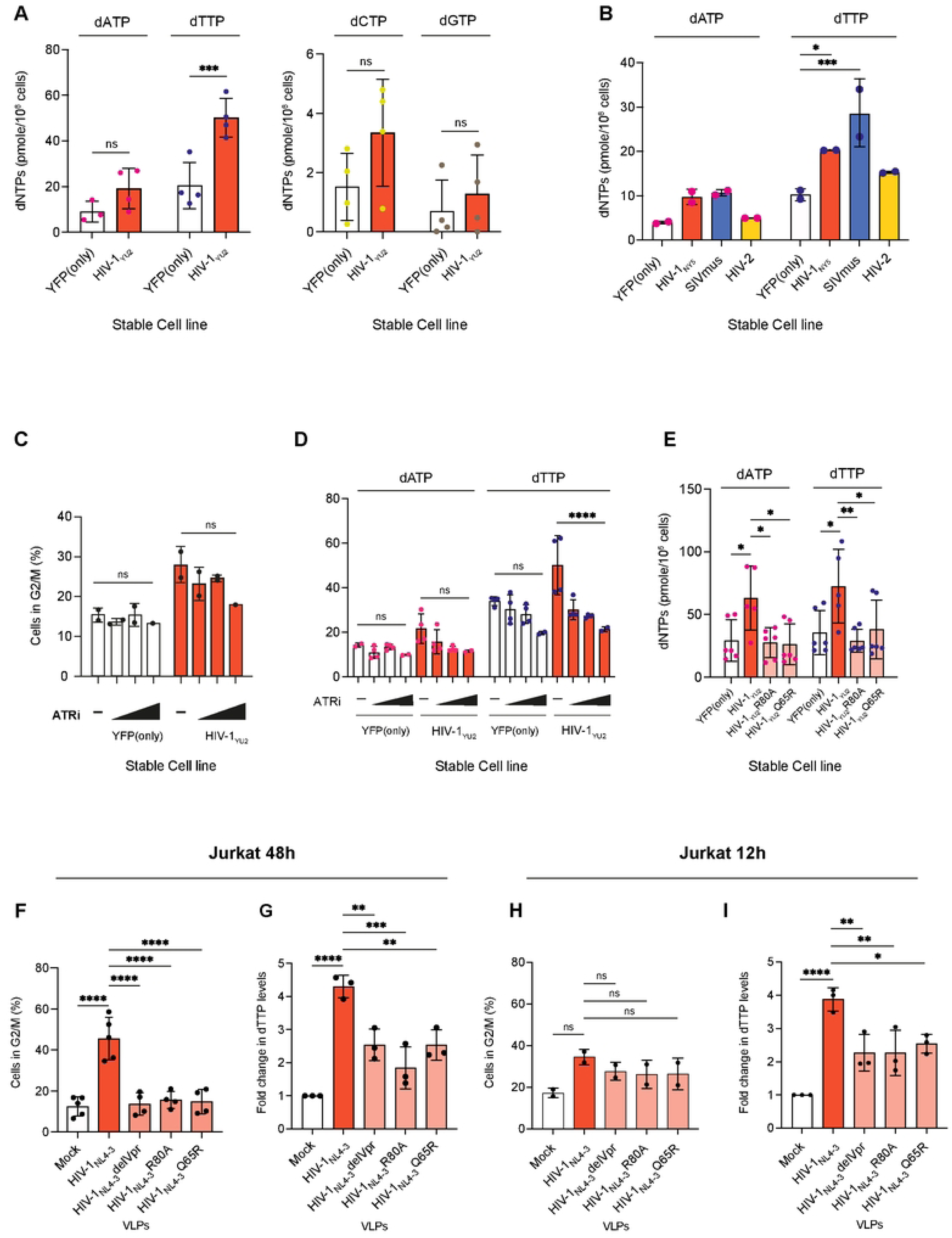
The effect of Vpr on dNTP levels. (A-D) HeLa stable cell lines encoding the indicated Vpr protein were induced with doxycycline for 48h. (C&D) Increasing concentration of the ATR inhibitor (VE-821) was added to cells for a further 24h. (A, B & D) YFP positive cells were sorted and the levels of individual dNTPs were determined by radioactive primer extension assay. (C) Cells were fixed, stained with DAPI and the percentage of cells in G2/M was analysed by flow cytometry. (E) Jurkat stable cell lines encoding HIV-1_YU2_, HIV-1_YU2_(R80A), HIV-1_YU2_(Q65R) or YFP(only) were induced with doxycycline for 48h. YFP positive cells were sorted and dATP/dTTP levels determined by primer extension assay. (A-E) Individual points represent separate biological repeats. Bars show the mean and SEM. Statistical analysis was performed using two-way ANOVA to calculate significant differences from the control YFP(only) or mock treated cells. The significance level was set at * p < 0.05, ** p < 0.01, *** p < 0.001,**** p < 0.0001. (F-I) Jurkat cells were infected with VLPs carrying WT or mutant Vpr as indicated or left uninfected (mock). (F) 48h post-infection and (H) 12h post-infection, cells were fixed, stained with DAPI and the percentage of GFP positive (infected) cells in G2/M was analysed by flow cytometry. (G) 48h post-infection and (I) 12h post-infection, live, GFP positive cells were sorted and levels of dTTP determined by primer extension assay. The fold change compared to uninfected cells is plotted. Individual points represent separate biological repeats. Bars show the mean and SEM of at least two biological repeats. Statistical analysis was performed using one-way ANOVA to calculate significant differences from the HIV-1_NL4-3_ treated cells. The significance level was set at * p < 0.05, ** p < 0.01, *** p < 0.001,**** p < 0.0001. ns indicates not significant.

### Virion-incorporated Vpr induces cell cycle arrest and increases levels of dNTPs within 12 hours

Vpr is actively packaged into lentiviral virions, implying an important role during early replication, before *de novo* synthesis of Vpr occurs in infected cells [7]. To determine if virion-associated Vpr was sufficient to cause inhibition of both G2 and M phases of the cell cycle and raise dNTP levels, we used the HIV-1 p8.2 *gag-pol* plasmid, that expresses Vpr and the other HIV-1 accessory proteins in addition to Gag-Pol, to generate HIV-1 virus-like particles (VLPs). These HIV-1 VLPs packaged Vpr but could not express nascent Vpr during infection as *vpr* is not encoded as a gene in the VLP genome and therefore no further expression of Vpr occurs after the initial protein is delivered. We also introduced mutations into the p8.2 plasmid to generate the HIV-1_NL4-3_(R80A) or HIV-1_NL4-3_(Q65R) mutants, or a stop codon at the start of Vpr (delVpr). First, we determined if virion-associated Vpr could induce both G2 and mitotic arrest. HeLa and Jurkat cells were infected with increasing amount of WT, R80A, Q65R or delVpr VLPs or left uninfected (mock). Two days post-infection, cells were analysed by flow cytometry for the percentage of GFP positive (VLP positive) cells in G2 and mitosis (**Fig.S6**). As seen with transduction of Vpr alone, virion-associated wild type HIV-1_NL4-3_ Vpr was able to increase the percentage of cells in both G2 (**Fig.S6A**) and M (**Fig.S6B**) in HeLa cells in a dose dependent manner but only arrested Jurkat cells in G2 not M (**Fig.7F and Fig.S6C and D**). VLPs carrying mutant HIV-1_NL4-3_ Vpr did not increase the percentage of cells in G2 or M in either cell line (**Fig.7F and Fig.S6**). This shows that the Vpr delivered to cells in virions is sufficient to induce the same cell cycle arrest phenotypes as Vpr expressed directly in cells. We then measured dNTPs levels in VLP infected cells two days post infection. As seen in the stable Vpr cell lines, virion-associated Vpr also induced a rise in dTTP levels (**Fig.7G**). As higher dNTP levels may facilitate reverse transcription and/or integration which typically occur within 12 hours of infection, we analysed the effect of Vpr on both the cell cycle and dNTP levels 12h post infection (**Fig.7H and I**). Excitingly, virion-incorporated Vpr was able to induce an increase in dTTP levels (**Fig.7I**) at this time point, despite there being only a small increase in cell cycle arrest (**Fig.7H**). Moreover, the increase in dTTP levels was similar to that seen 48h post infection (**Fig.7G**), implying that no further increase occurred with time. Interestingly, when we enriched uninfected populations of HeLa and Jurkat T-cells for cells in G2, by collecting the larger cells from the population by flow cytometry, and analysed the dTTP levels in these cells, we found that dTTP levels were increased by less than 2-fold compared to the unsorted cells (**Fig.S6F**), as opposed to the 4-fold increase caused by Vpr (**Fig.7G and 7I**), despite more cells being in G2 than in our Vpr-infected cells (**Fig.S6E**). This implies that Vpr induces a larger increase in dNTP levels without this leading to increased cell cycle arrest.

## Discussion

It is well documented that Vpr causes cell cycle arrest at the G2/M boundary and this has been reproducibly linked to ATR activation [13, 15, 42]. However, the involvement of ATM is more controversial and the mechanism of activation of DNA damage pathways and how the known targets of Vpr are connected to the cell cycle block is unknown. Vpr has been shown to induce BRCA1 and γH2AX foci [13], but there is disagreement about whether Vpr actually induces DNA damage, particularly double strand breaks (DSB) [15, 48–52], or if it instead activates the DNA replication checkpoint. Indeed, it has been proposed that Vpr activates the G2/M checkpoint through an unusual replication stress pathway that initiates in S-phase, through activation of Chk1 via Ser345 phosphorylation [53]. Many of the discrepancies likely arise from the systems used to analyse the effect of Vpr.

Here, we confirm that the G2 arrest requires ATR but not ATM (**Fig.5**), and that this activity is conserved across Vpr proteins from many different SIV strains [48, 54]. However, additionally, we show that some Vpr proteins, namely from HIV-1, SIV_cpz_ and HIV-2, also cause a further block in the M phase of the cell cycle (**Figs 1**, **2 and 3**) and this block requires both ATR and ATM (**Fig.5**). We believe that as there are generally far fewer cells in the population in M-phase at any one time, this block has previously been overlooked. Furthermore, most studies have only used DAPI staining in flow cytometry to look at which phase of the cell cycle cells are in, which cannot separate whether cells are in G2 or M phase. ATR is primarily activated in response to DNA damage during S/G2, therefore its role in mitotic arrest may be an indirect consequence of inhibiting G2 arrest upstream. However, it has also been shown to function in faithful chromosome segregation during mitosis, and thus may be directly involved in Vpr-induced mitotic arrest [55]. Our immunofluorescence experiments revealed that HIV-1 Vpr causes multipolar spindles in mitosis, and this is accompanied by γH2AX accumulation. Although Vpr reportedly relocalised PLK-1 in fission yeast [17], the chromosomal arrangements in Vpr positive cells here did not resemble those in cells treated with a PLK-1 inhibitor, BI2536 (**Fig.S4B**), suggesting that the mechanism may not be the same in mammalian cells. Nonetheless, the amalgamation of G2 and M populations in most analysis and the requirement for ATM in mitotic arrest but not G2 arrest could help explain conflicting reports about ATM function in Vpr-induced cell cycle arrest [13, 16].

Using Vpr mutants, we further showed that, like the G2 arrest, M arrest is also dependent on DCAF binding and residues in the CtD of Vpr. It is possible that the cells stuck in M phase may just be cells that have evaded the G2/M checkpoint. However, the fact that not all Vpr proteins that inhibit cells in G2 also inhibit cells in M (**Figs 1**, **2 and 3**), and the fact that chimeric proteins between HIV-1 and SIV_mus_ Vpr proteins appear to be able to induce an M-phase block without inhibiting G2 (**Fig.4**) suggests that these are distinct blocks.

Interestingly, SIV_mus_ Vpr did not cause such a strong block to G2 as HIV-1 Vpr in a single round of the cell cycle (**Fig.3**). However, the number of cells in G2 increased over 72 hours suggesting that cells were blocked in subsequent cycles and accumulated in G2. This did not appear to happen in the presence of HIV-1 Vpr where the percentage of cells in G2 remained constant and the percentage of cells in M phase actually reduced. For HIV-1 Vpr, the initial G2 block was very strong and with the additional block in M phase, overall, less cells were continuing to cycle. We also observed high levels of cell death following prolonged HIV-1 Vpr expression which removed cells from the analysis. Therefore, although the G2 block is more conserved, Vpr proteins from different lentiviruses induce G2 arrest with different kinetics, suggesting possible differences between species. Some variation may be explained by the fact that we could not use the cognate cells to match each Vpr strain. Some Vpr proteins may have a stronger effect in the simian species in which they evolved. This appears to be the case with SIV_agm_ Vpr, which induces G2 arrest in AGM cells but not HeLa cells (**Fig.2**) [32, 54, 56]. It is possible that some additional Vpr proteins will inhibit mitosis in their cognate cells.

Although we cannot discount other forms of DNA damage caused by Vpr, we have shown that HIV-1 Vpr induces ultra-fine anaphase bridges (UFB) (**Fig.6**). This helps to explain the role of MUS81 in Vpr function. MUS81 was initially described as being modulated through the interaction of Vpr and SLX4 and being linked to G2 arrest [22, 54]. However, the requirement for SLX4 has since been disputed and MUS81 degradation is genetically separable from G2 arrest [48, 57, 58] (**Fig.6**). MUS81 is important for genome integrity. It is required for processing of UFB that can be formed either by collapsed replication forks during S phase or by unresolved homologous recombination (HR) intermediates at mitosis [43]. Ongoing replication stress can be marked by FANC2D foci, which has been shown to accumulate in the presence of Vpr [22]. However, this usually leads to cell cycle arrest before mitosis. Here, we show that ectopic expression of MUS81 relieves the mitotic block induced by HIV-1 Vpr without affecting G2 arrest, implying that MUS81 is only necessary for mitotic arrest. (**Fig.6**) Moreover, we can detect UFB during mitosis in these cells, suggesting that Vpr expression induces UFB via homologous recombination intermediates. Intriguingly, MUS81 was also degraded by SIV_mus_ and HIV-1_YU2_R80A that do not induce mitotic arrest, indicating that degradation of MUS81 is not sufficient to generate UFB or M arrest, as well as being disconnected from G2 arrest [57, 58]. Importantly, SIV_mus_ and HIV-1_YU2_R80A Vpr proteins do not induce UFBs (**Fig.6**) so there is nothing for MUS81 to resolve. As it is not required in this case, it is perhaps not surprising that degrading it does not affect cell replication. Overall, degradation of MUS81 is necessary but not sufficient to arrest cells in mitosis. The conservation of MUS81 degradation implies that another of its functions hinders lentiviral replication more generally. Pertinently, the accumulation of UFB did correlate with M arrest. Overall, this suggests a two-step mechanism for induction of M arrest, as suggested by Li et al. [49]. First, Vpr must induce DNA damage which leads to the formation of UFB during HR. Second, Vpr must also induce the degradation of the host cell machinery that resolves such UFB. This then activates the spindle assembly check point and the cell cycle is paused. Whether the same damage leads to alternative DDR pathways that activate ATR and result in G2 arrest remain to be seen. Interestingly, DNA damage and activation of DDR pathways was also found to be independent of repression of HR and G2 arrest using a panel of Vpr mutants and variants [49]. Other reported targets of Vpr are involved in chromatin remodelling and gene silencing [26, 51] and it is possible that modulation of these factors alters chromatin structure and triggers DNA damage. Indeed, stimulating the decondensing of chromatin would likely increase the number of potential integration sites and thus enhance infection, particularly in non-dividing cells, which is a noted effect of Vpr.

Despite speculation, the biological significance of Vpr-induced cell cycle arrest is still unclear, particularly as Vpr appears to have a greater effect on HIV-1 replication in non-dividing cells. It has been proposed that Vpr is required to suppress human immune function by preventing T-cell clone expansion [59] or blocking innate immune sensing [22, 60]. Alternatively, an initial hypothesis was that HIV-1 expression was optimal in G2, and so Vpr activity was creating a favourable cellular environment for viral replication [61]. Here, we have shown that Vpr-induced G2 arrest correlates with an increase in dNTP levels. Interestingly, Vpr induces a greater fold increase in dNTP levels than is normally observed in the G2 phase of the cell cycle (**Fig.7 and Fig.S6**). Importantly, the rise in dNTP levels happens early in infection, and before expression of nascent viral proteins, indicating that the Vpr protein packaged into viral particles is sufficient to generate this effect. By increasing dNTP levels, Vpr expression has a similar phenotype to Vpx expression, albeit via a different mechanism [62]. We feel this is likely a beneficial side effect of Vpr activity as DNA damage has been reported to promote nucleotide biosynthesis [63]. It is intriguing that some Vpr proteins can also target SAMHD1 for degradation, raising the possibility that this function arose to augment the favourable conditions induced by the parental Vpr. Interestingly, the papillomavirus HPV31 has been reported to regulate the levels of RRM2, a component of the ribonucleotide reductase complex that generates *de novo* dNTPs, resulting in increased dNTPs that are essential for replication [64]. RRM2 levels are enhanced through the viral E7 protein and activation of the ATR-Chk1 axis of the DNA damage response. Normally, RRM2 is degraded in G2 via the Skp1-Cul1-F-box complex in combination with cyclin F to maintain balanced dNTP pools. After DNA damage, cyclin F is downregulated in an ATR-dependent manner to allow accumulation of RRM2, and thereby dNTPs that are necessary for DNA repair [65]. In our experiments, we saw that dNTP levels were reduced to control levels when an ATR inhibitor was added to infected cells, and that this reduction in dNTPs occurred at lower concentrations of inhibitors than the level required to inhibit the G2 block. This suggests that Vpr increases dNTP levels by virtue of activating ATR rather than as a consequence of arresting the cell cycle. It also provides an alternative advantageous outcome of ATR activation by Vpr to the proposal that ATR activation is linked to evasion of innate immune sensing [22].

In summary, Vpr proteins from a broad range of primate lentiviruses degrade cellular proteins involved in repairing DNA damage, including MUS81, although none of the known targets appear to be responsible for the G2 arrest observed. Presumably, either the degradation of a combination of these factors or a yet-to-be-identified factor induces ATR activation, and this starts a cascade that results in increased dNTP levels and a cell cycle block. Additionally, some Vpr proteins, including HIV-1 Vpr, induce DNA damage leading to the formation of UFB in mitosis that cannot be resolved, activating ATM and the SAC. Together, these activities result in cell cycle blocks at G2 and M. The rapid activation of ATR and ATM suggest that either the DNA damage that triggers their activation or the kinases themselves enhances the early stages of replication, most likely integration. However, it is interesting that the last stage of integration actually requires DNA repair and so these different activities must be finely balanced. Clearly, understanding the mechanism of Vpr-induced cell cycle arrest may provide insights into the requirements for efficient HIV-1 infection.

## Materials and methods

### Cell lines

HEK293T, HeLa, U2OS and CV-1 cell lines were maintained in Dulbecco’s modified Eagle medium (DMEM, Thermo Fisher), and Jurkat and SupT1 T-cells were incubated in RPMI-1640 (Thermo Fisher). All cell lines were from Bishop laboratory cell stocks authenticated by short-tandem repeat (STR) profiling and tested mycoplasma-free. Media was supplemented with 10% heat-inactivated foetal bovine serum (FBS; Biosera) and 1% Penicillin/Streptomycin (Sigma). Cells were grown in a humidified incubator at 37°C and 5% CO_2_.

### Plasmids and primers

The plasmids used to produce virus-like particles (VLP), KB4 (MLV *gag-pol*), pCMVΔ8.91 (HIV-1_NL4-3_ *gag-pol*), pCMVΔ8.2 (HIV-1_NL4-3_ *gag-pol*), pVSV-G and pCSGW, have been described before [66–69]. pCMVΔ8.2delVpr was obtained from the HIV reagent program.

For transduction of Vpr, FLAG-HA-SIV_deb_ Vpr was initially synthesised by GeneArt (Thermo Fisher) and cloned into our pLRGatewayIRESeYFP plasmid [46] using BamHI (into a BglII site) and XhoI restriction enzymes. The remaining Vpr genes and the HIV-1/SIV_mus_ Vpr chimeras were synthesised untagged by GeneArt (Thermo Fisher) and cloned into pLRGatewayIRESeYFP-FLAG-HA-Vpr_deb_ using BglII and XhoI restriction sites (see Table 1 for Vpr sequences). C-terminal truncations were amplified with primers listed in Table 2 and cloned into pLRGatewayIRESeYFP-FLAG-HA-Vpr_deb_ using BglII and XhoI restriction sites. Mutations in the Vpr gene of pCMVΔ8.2 (Q65R, R80A and H71R) or pLRGatewayIRESeYFP-Vpr plasmids were synthesised using the QuickChange II-XL site-directed mutagenesis kit (Agilent) according to manufacturer’s instructions and primers listed in Table 2. To generate the pLRGatewayIRESmApple plasmids, mApple was cloned in place of the eYFP gene using the NEBuilder HiFi DNA Assembly Cloning Kit (NEB) and BstXI and NotI restriction sites as per the manufacturer’s instructions. To make stable cells lines expressing Tet-inducible Vpr, individual FLAG-HA tagged Vpr genes were amplified from the pLRGatewayIRESeYFP-Vpr plasmids using the primers in Table 2 and subcloned into pLVX-TetOne-puro (Clontech) using EcoRI and BamHI restriction sites. The IRES-EYFP fragment was cloned into pLVX-TetOne-puro to make pLVX-TetOne-IRES-eYFP-Puro.

**Table 1:**
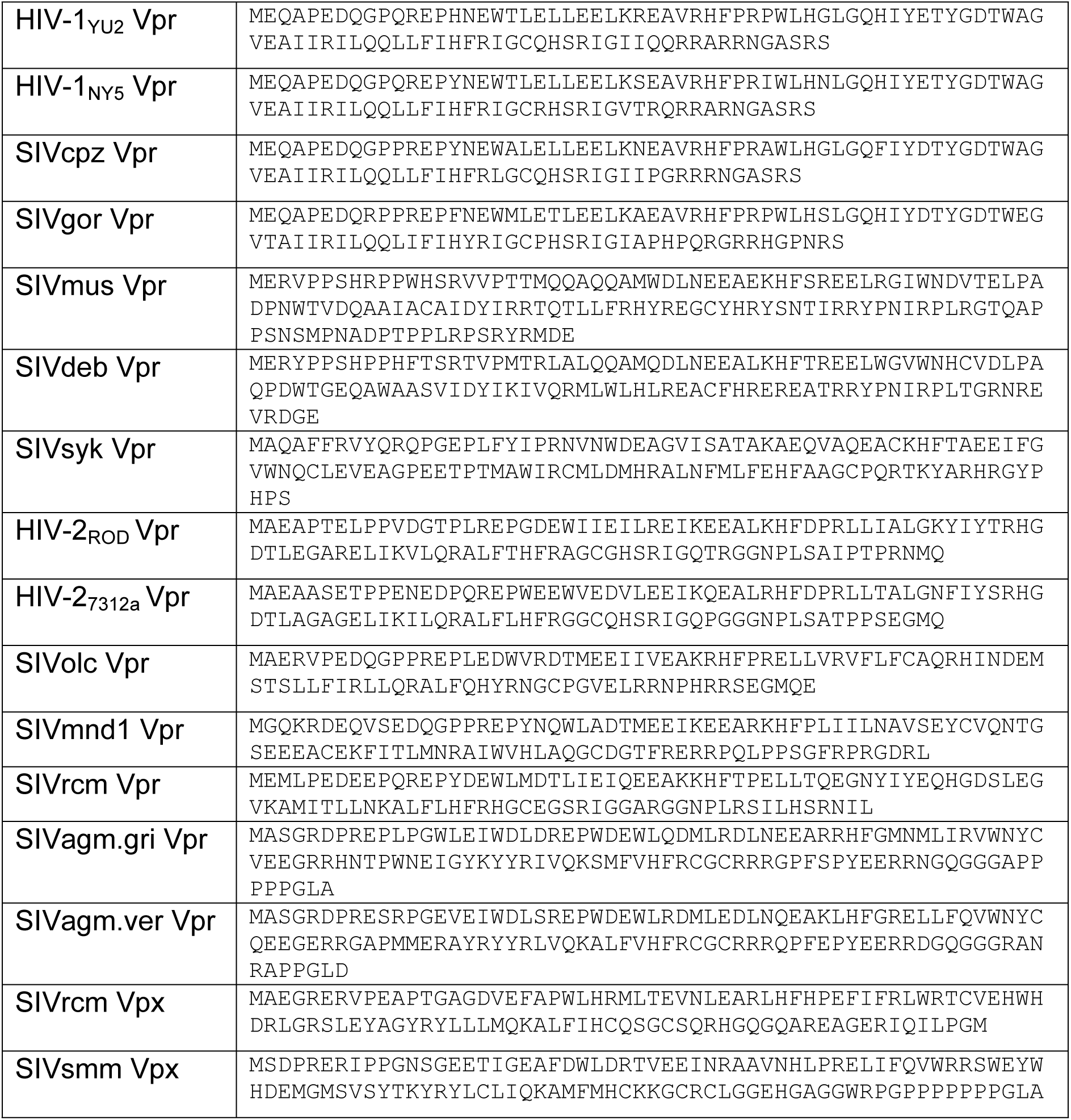
Vpr and Vpx protein sequences.

**Table 2:**
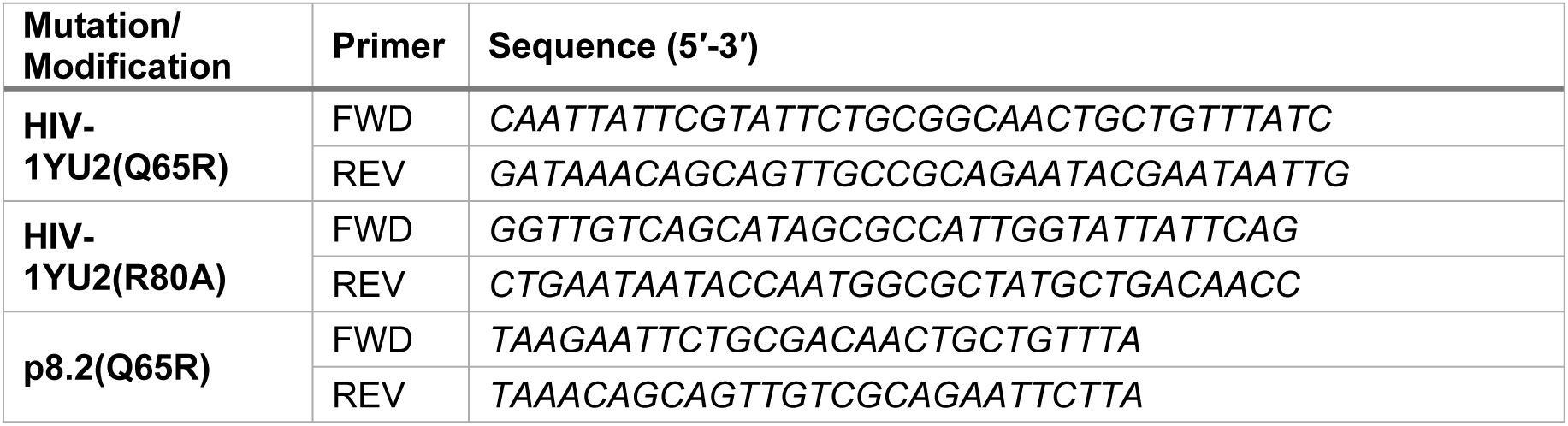
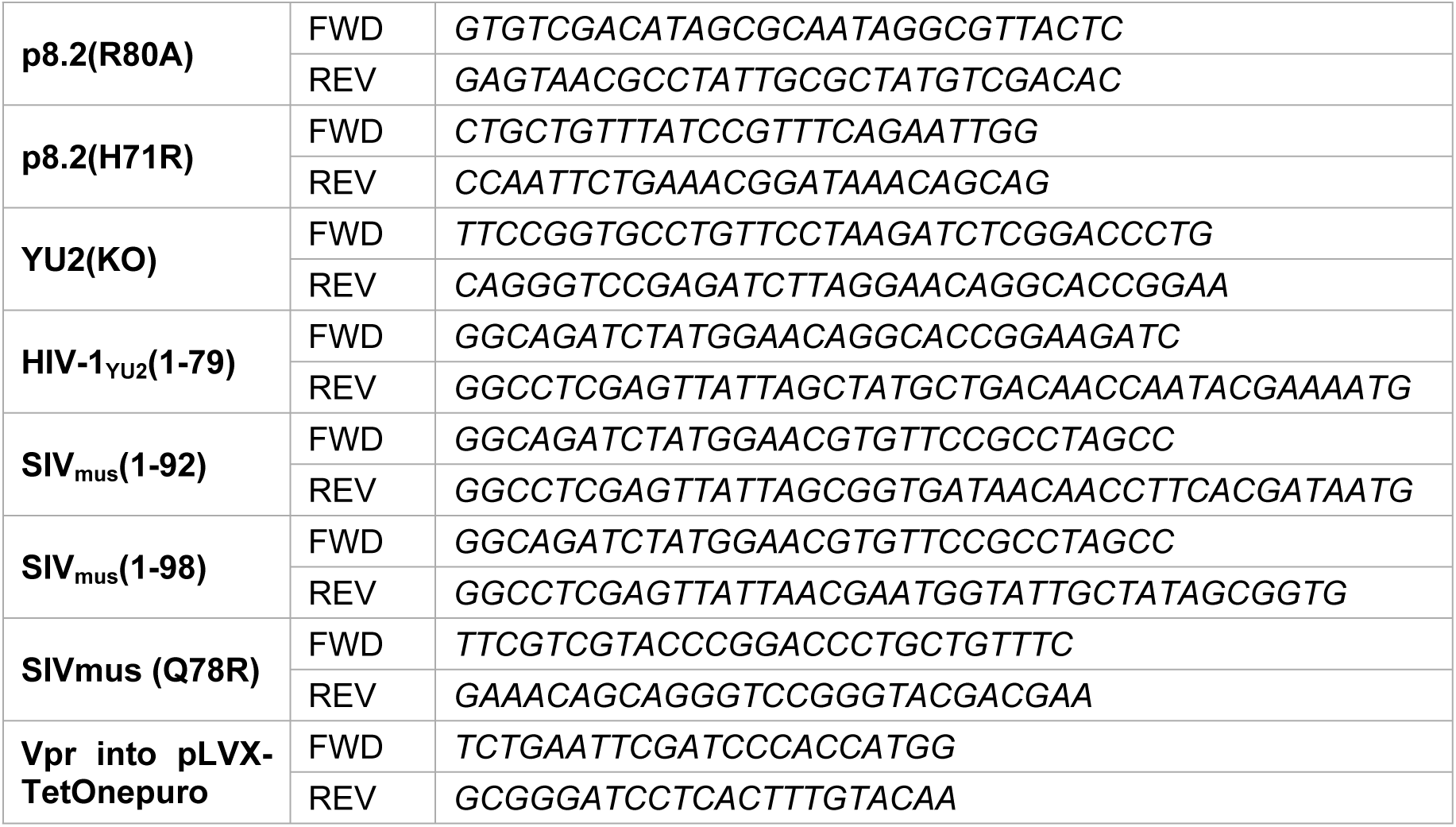
Primers used for cloning.

To generate the CCDC137 expression vector, Myc-CCDC137 was synthesised by GeneArt and cloned into pLRGatewayIRESmApple using the EcoNI and XhoI restriction sites. Expression vectors, pCDNA3.1-GFP-MUS81, pCDNA3.1-GFP(only) and XPF-GFP were kindly gifted from Stephen West.

### Vector/ Virus-like-particle production

To generate MLV vectors encoding Vpr or Vpx, HEK293T cells were co-transfected with pKB4, pVSV-G and either pLRGatewayIRESeFYP-Vpr/Vpx or pLRGatewayIRESmApple-Vpr/Vpx at a ratio 1:1:1 with TransIT-lenti (Mirus, MIR 6603). To generate HIV vectors to make stable cell lines, HEK293T cells were co-transfected with pCMVΔ8.91, pVSV-G and pLVX-TetOne-IRES-eYFP-Puro-Vpr at a ratio 1:1:1 with polyethylenimine (PEI [70]). To generate HIV-1 VLP carrying Vpr proteins, HEK293T cells were co-transfected with pCMVΔ8.2, pVSV-G and pCSGW at a ratio of 10:1:2.6 with PEI. Approximately 16h post-transfection, cells were treated with 10mM sodium butyrate for 8h and VLP-containing supernatants were harvested 16h later. To measure viral titres, HeLa cells were transduced with serially diluted supernatant containing the VLPs. After 72h, cells were harvested and percentage of YFP or mApple positive cells determined by flow cytometry.

### Stable Cell Lines

To generate the stable cell lines, HeLa cells were transduced with a high volume of HIV-1 vectors encoding FLAG-HA-Vpr-IRES-YFP downstream of a doxycycline-inducible TET-One 3G (TET-3G) promoter flanked by HIV-1 LTRs. After 72h, doxycycline (1 μg/ml) was added for 24h. YFP positive cells were then sorted using an Aria II influx cell sorter, re-plated and grown up for use in assays.

### Edu Assay

HeLa stable cell lines were induced with doxycycline (1 μg/ml) for 24h before incubation with complete media containing Edu (10 μM) and doxycycline (1 ug/ml) for 30 minutes. Cells were washed twice with complete media and then either harvested (0h timepoint) or further incubated with complete media containing doxycycline (1 μg/ml). Harvested cells were fixed with 4% paraformaldehyde (J60401.AK, Thermo fisher). Edu was activated using the CLICK-iT Edu Kit (Invitrogen, C10646) as per the manufacturer’s instructions and cells were immunolabelled with anti-MPM2-Cy5 (EMD Millipore, 16-220). YFP and Edu positive cells were analysed by flow cytometry. Cells were initially separated into early (S1), middle (S2), and late (S3) S phase (**Fig.S2F**). At 0h, these subpopulations represent S phase as Edu is only incorporated during DNA synthesis. At later time points, Edu positive cells begin to transition through the cell cycle. Therefore, cells in S1 represent the G1/early S population while S3 represent the late S/G2/M population. S2 represents cells in S phase only. The S3 subpopulation was further separated into MPM2 positive cells to determine the percentage of mitotic cells. A representative FACS plot outlining the gating strategy is shown (**Fig.S2F**). The percentage of cells in G1/early S, S phase, Late S/G2/M was determined. The percentage of cells in M phase was further calculated and removed from the Late S/G2/M population to calculate the Late S/G2 population size.

### Infection/transduction

Cells were infected with equal numbers of HIV-1 VLP, normalised by TAQ-pert assay [71] in complete media containing 3 μg/ml polybrene, via spinoculation for 90 minutes at 1,000 rpm at 15°C. Cells were then incubated at 37°C for the remainder of the experiment.

### Flow cytometry

Cells were washed once with PBS and then fixed in ice cold 4% paraformaldehyde (J60401.AK, Thermo fisher) for 10 minutes at room temperature. Cells were then washed, permeabilised in 0.1% triton X-100 (FC005, R&D systems), washed again and labelled with DAPI (1 μg/ml). Live cells were gated and analysed on either a LSRFortessa or Fortessa X-20 analysers (BD Biosciences). Data was analysed using FlowJo software. For MUS81 degradation assay, the GFP median fluorescent intensity (MFI) of the mApple positive population was determined. For cell cycle analysis, cells were incubated with the anti-MPM2-Cy5 (EMD Millipore, 16-220), diluted in PBS (1:200). The percentage of YFP-positive cells in G1/S, G2 and M was determined. For DNA damage analysis, cells were labelled with anti-H2AX(p)-PE (CST, 5763S) and anti-MPM2-Cy5 (EMD Millipore, 16-220). The percentage of phosphorylated H2AX (γH2AX) cells was determined in each cell cycle phase.

### Cell sorting

Cells were washed in ice cold PBS and resuspended in PBS containing 2% FCS. Cells were either treated with 7AAD (Biorad, 1351102) or left unstained. Single cell populations were sorted using the Avalon S3e into PBS. For the dNTP assay, cells cells were immediately centrifuged, supernatant removed and frozen on dry ice, before storing in -80°C.

### Immunoblotting

Cells were lysed in ice cold radioimmunoprecipitation assay (RIPA) buffer (Thermo Fisher) containing protease inhibitor cocktail (Roche) and DNAse (Thermo Fisher, #88701). Proteins were separated on 4-12% Bis-Tris SDS-PAGE gels (Thermo Fisher) and transferred to a PVDF membrane for immunoblotting. Primary antibodies used: anti-HA (C29F4, CST), anti-GFP (Santa Cruz, sc-9996), anti-tubulin-α (Bio-Rad, VMA00051), anti-MUS81 (Abcam, ab14387). Secondary antibodies used: anti-rat IRDye 680CW (LI-COR, 926-68076), anti-mouse IRDye 680RD/800CW (LI-COR, 926-68072/926-32212) and anti-rabbit IRDye 700CW/800CW (LI-COR, 926-68073/926-32213). Blots were visualised on an Odyssey Clx imaging system (LI-COR) and protein expression quantified using the LI-COR Image studio 5.5.4 software. GFP-MUS81 levels were normalised against tubulin-α.

### Immunofluorescence

HEK29T were grown on glass coverslips and transfected with pLRGateway-Vpr-IRES-eYFP plasmids. For spindle visualisation, untransfected cells were treated with either Nocodazole (M1404, Merck, 50 µg/ml) or BI2536 (#26744, Cell Signaling Technology, 100 nM) for 15h. For visualising ultra-fine DNA bridges, cells were washed with PTEMF buffer (20 mM PIPES pH 6.8, 0.2% Triton X-100, 1 mM MgCl_2_ 10 mM EGTA and 4% paraformaldehyde) for 10 minutes with agitation. For all other IF, cells were fixed in 4% paraformaldehyde for 10 minutes at room temperature. Cells were then permeabilised with 0.1% triton X-100 and blocked in 3% BSA in PBS. Cells were labelled with the following primary antibodies: anti-RPA (ab2175, ABCAM, 1:1000), anti-FANCD2 (NB100-182, Novus, 1:1000), anti-YFP (ab6556, abcam, 1:2000), anti-CycB (sc-245, Santa-Cruz, 1:300) or anti-tubulin (T5168, Merck, 1:40000) and the following secondary antibodies: Rabbit Alexa Fluor 647 (Life Technologies, a31573), Mouse Alexa Fluor 568 (Abcam, ab175472). Coverslips were mounted on glass slides with ProLong Gold Antifade Mountant with DAPI (Thermo Fisher, 15260719). Cells were imaged with the Leica SP5 100X 1.3NA oil immersion objective (Leica). The number of mitotic cells undergoing anaphase with ultra-fine bridges was determined blind.

### Inhibitors

To induce cell cycle arrest, cells were treated with nocodozale (Merck, M1404) for 16h diluted to 1:10000 or RO-3306 (Sigma, SML0569) for 24h at a concertation at 9 μM. To induce DNA damage, cells were treated with hydroxyurea (HU) at 200 μM for 8h. To inhibit the spindle assembly checkpoint cells were treated with Go6976 for 1h at a concentration of 2 μM. To inhibit ATR or ATM, cell were treated with VE-821 (ATRi) (Sigma, SML1415) or KU-55933 (ATMi) (Sigma, SML1109) at 10 μM for 24h, respectively. As a control, cells were treated with DMSO only.

### dNTP quantification

Levels of dATP, dCTP, dGTP and dTTP were measured, as previously described [9, 46], in HeLa or Jurkat cells infected with HIV-1 VLP carrying wild type or mutant Vpr for the indicated time, or in stable HeLa or Jurkat Vpr cell lines induced with doxycycline for 48h. YFP positive cells were sorted on a S3e cell sorter (Bio-Rad). For dATP and dTTP measurements, 2 × 10^5^ cells were analysed. For dCTP and dGTP, 2 × 10^6^ and 4 × 10^6^ cells were analysed respectively. A proportion of cells were removed to confirm cell cycle arrest. ATRi was added to cells 24h before harvesting, at either 1, 2.5 or 5 μM. dNTP levels were quantified by radiolabel incorporation assays, performed using the following Duplex Extension Templates (Merck): Common Hot start Primer: 5’-CGTGCACCGCCTCCACCGCC-3’; Bottom strand dATP Extension Hot start Long version 2: 5’- CCCTCCCTCCCTCCCTCCCTGGCGGTGGAGGCGGTGCACG-3’; Bottom strand dCTP Extension Hot start Long version 2: 5’- CCCGCCCGCCCGCCCGCCCGGGCGGTGGAGGCGGTGCACG-3’; Bottom strand dGTP Extension Hot start Long version 2: 5’- AAACAAACAAACAAACAAACGGCGGTGGAGGCGGTGCACG-3’; Bottom strand TTP Extension Hot start Long version 2: 5’- CCCACCCACCCACCCACCCAGGCGGTGGAGGCGGTGCACG-3’.

## Quantification and statistical analysis

Statistical analysis of data was performed using GraphPad Prism 9 software and indicated in the figure legends. Significance is reported with one, two, three or four asterisks if p < 0.05, p < 0.01, p < 0.001 or p < 0.0001, respectively. NS = not significant. Separate biological repeats are plotted as individual points for each experiment to allow readers to observe the data range directly.

## Acknowledgements

We thank Jonathan Stoye and Bart Szafran for helpful discussions. For the purpose of Open Access, the authors have applied a CC BY public copyright licence to any Author Accepted Manuscript version arising from this submission.

## Supporting Information

S1 Fig. Vpr panel protein expression

S2 Fig. Statistical analysis of data in Fig 2

S3 Fig. Synthesis of stable, inducible Vpr cell lines

S4 Fig. The spindle assembly checkpoint and Vpr

S5 Fig. The effect of overexpressing potential Vpr target proteins on cell cycle arrest

S6 Fig. The effect of virion-incorporated Vpr on cell cycle arrest

## Notes

### Competing Interest Statement

All authors, except S.J.B., declare no competing interests. S.J.B. is a co-founder, VP Science Strategy and a shareholder of Artios Pharma Ltd.

